# Identification and receptor mechanism of TIR-catalyzed small molecules in plant immunity

**DOI:** 10.1101/2022.04.01.486681

**Authors:** Shijia Huang, Aolin Jia, Wen Song, Giuliana Hessler, Yonggang Meng, Yue Sun, Lina Xu, Henriette Laessle, Jan Jirschitzka, Shoucai Ma, Yu Xiao, Dongli Yu, Jiao Hou, Ruiqi Liu, Huanhuan Sun, Xiaohui Liu, Zhifu Han, Junbiao Chang, Jane E. Parker, Jijie Chai

**Author notes:** Equal contribution authors (Shijia Huang, Aolin Jia, Wen Song, Giuliana Hessler, Yonggang Meng).

## Abstract

Plant nucleotide-binding leucine-rich-repeat receptors (NLRs) with an N-terminal toll/interleukin-1 receptor (TIR) domain sense pathogen effectors to enable TIR-encoded NADase activity for immune signaling. TIR-NLR (TNL) signaling requires conserved helper NLRs NRG1 and ADR1 and the lipase-like protein EDS1 that functions as a heterodimer with each of its paralogs PAD4 and SAG101. We show that TIR-containing proteins catalyze production of 2’-(5’’-phosphoribosyl)-5’-adenosine mono-/di-phosphate (pRib-AMP/ADP) *in vitro* and *in planta*. Biochemical and structural data demonstrate that EDS1-PAD4 is a receptor complex for pRib-AMP/ADP. pRib-ADP binding triggers a conformational change in the PAD4 C-terminal domain to allosterically promote EDS1-PAD4 interaction with ADR1-L1 but not NRG1A. Our study identifies TIR-catalyzed pRib-AMP/ADP as a missing link in TIR signaling via EDS1-PAD4 and as likely second messengers for plant immunity.

---

Nucleotide-binding leucine-rich repeat (NLR) proteins are intracellular immune receptors with crucial roles in innate immunity of plants and animals (*1*). Plant NLRs have a conserved organization with a central nucleotide-binding domain (NBD), a C-terminal leucine-rich repeat (LRR) domain and an N-terminal coiled-coil (CC) or Toll/interleukin-1 receptor domain, referred to as CNL or TNL receptors, respectively. Monocot plants lack TNLs but encode TIR-only and TIR-NB proteins (*2, 3*). Diversified plant NLRs mediate specific recognition of pathogen effectors to initiate effector-triggered immunity (ETI). Activated ETI signaling results in pathogen restriction and often a host hypersensitive cell death response (HR) at infection sites (*4*). Direct or indirect recognition of effectors induces NLR oligomerization (*5*), resulting in formation of large protein complexes termed resistosomes (*1, 6, 7*). The *Arabidopsis* CNL ZAR1 resistosome functions as a calcium-permeable channel for ETI signaling (*8*). By contrast, TNL resistosomes *Arabidopsis* RPP1 (*9*) and *Nicotiana benthamiana* (*Nb* tobacco) Roq1 (*10*) are NADase enzymes encoded by their N-terminal TIR domain (*11, 12*). TIR NADase activity relies on the presence of a conserved glutamate residue that is also essential for TIR-triggered immune signaling (*11, 12*).

Plant TIR-dependent immune signaling requires downstream conserved helper NLRs (hNLRs) of the CNL class and the lipase-like protein family members EDS1 (Enhanced Disease Susceptibility 1), PAD4 (Phytoalexin Deficient 4) and SAG101 (Senescence-Associated Gene 101) (*13, 14*). The hNLRs divide into ADR1 (ADR1, ADR1-like 1,2 (ADR1-L1, 2)) and NRG1 (NRG1A, B, C) sub-families in *Arabidopsis*. EDS1 exclusive heterodimers with PAD4 and SAG101 confer basal immunity and ETI responses (*15–17*). The C-terminal ‘EP’ domain (shared by EDS1, PAD4 and SAG101) contains positively charged residues required for TNL signaling (*16, 18–20*). Phylogenomic (*16, 21*) and genetic (*16, 17, 19, 22-25*) studies suggest a functional partnership of EDS1-PAD4 with ADR1 and EDS1-SAG101 with NRG1. This is supported by induced associations of *Arabidopsis* EDS1-SAG101 and EDS1-PAD4 with NRG1 and ADR1, respectively (*19, 26*), that are required for pathogen resistance and potentially Ca^2+^-channel activity of hNLRs (*19, 27*). In a current model, products generated by the TIR-encoded NADase activity are intercepted by EDS1 heterodimers for activation of hNLRs (*19, 26*). Variant-cyclic ADP-ribose (v-cADPR) is a TIR-generated product (*11, 12, 28*), although its molecular identity and role in the plant immune system remain elusive. Thus, TIR-catalyzed chemical products mediating immune signaling are unknown.

Here, we identify plant TIR-catalyzed pRib-ADP and pRib-AMP through mass spectrometry (MS) and structural biology. These two compounds are produced by expression of the RPP1 resistosome or TIR domain of *Arabidopsis* RPS4 in insect cells. They exhibit binding to and promotion of EDS1-PAD4 direct interaction with ADR1-L1 but not NRG1A. The same activities are reproduced with chemically synthesized pRib-ADP and pRib-AMP. These molecules are much less efficient in promoting EDS1-SAG101 interaction with NRG1A, indicating preferential recognition of pRib-ADP and pRib-AMP by EDS1-PAD4. Structural comparisons reveal that pRib-ADP binding results in a conformational change in the EP domain of PAD4, enabling interaction with ADR1-L1. The data establish two nucleotide derivatives, pRib-ADP and pRib-AMP, as TIR-produced signals activating the EDS1/PAD4/ADR1 immunity module in plants. Identification of pRib-ADP and pRib-AMP as potential immune second messengers opens the way for designing small molecules to manipulate plant immunity.

## Reconstitution of TIR-induced EDS1-PAD4 interaction with ADR1 in insect cells

A TNL-triggered *Arabidopsis* EDS1/SAG101/NRG1 immunity signaling module was reconstituted in *Nb* tobacco (*16, 19*). To facilitate identification of TIR domain catalytic products, we tested whether NRG1 or ADR1 activation can be reconstituted in heterologous cells. We co-expressed *Arabidopsis* RPP1, its activating pathogen effector ATR1, EDS1, PAD4 (collectively called RAEP) and ADR1-L1 in insect cells and assayed for induced EDS1-PAD4 interaction with ADR1-L1 using a pull-down assay (*16, 26*). As shown in (Fig. 1A, top), EDS1-PAD4 but not EDS1-SAG101 co-purified with ADR1-L1 in the assay, supporting specific and direct interaction of EDS1-PAD4 with ADR1-L1. The interaction was dependent on presence of the RPP1 resistosome. EDS1-SAG101, but not EDS1-PAD4 interacted with NRG1A when co-expressed with RPP1 and ATR1 (called RAES) (Fig. 1A, bottom). These interactions were abolished when using an NADase catalytic mutant (E158A) of RPP1 (*9*) (Fig. 1A), suggesting that RPP1 TIR catalytic products promote EDS1 heterodimer associations with specific hNLR sub-types. The isolated TIR domain (residues 1-236) of *Arabidopsis* TNL RPS4 triggers *EDS1*-dependent HR cell death when expressed in *N. tabacum* plants (*29*). Co-expression of RPS4 TIR also promoted EDS1-PAD4 and EDS1-SAG101 binding to ADR1-L1 and NRG1A, respectively, which was abolished by mutation of the RPS4 TIR catalytic residue Glu88 (Fig. 1B). These results show that TIR domain catalytic products induce EDS1-PAD4 and EDS1-SAG101 interactions with their co-functioning hNLRs.

**Fig. 1.**
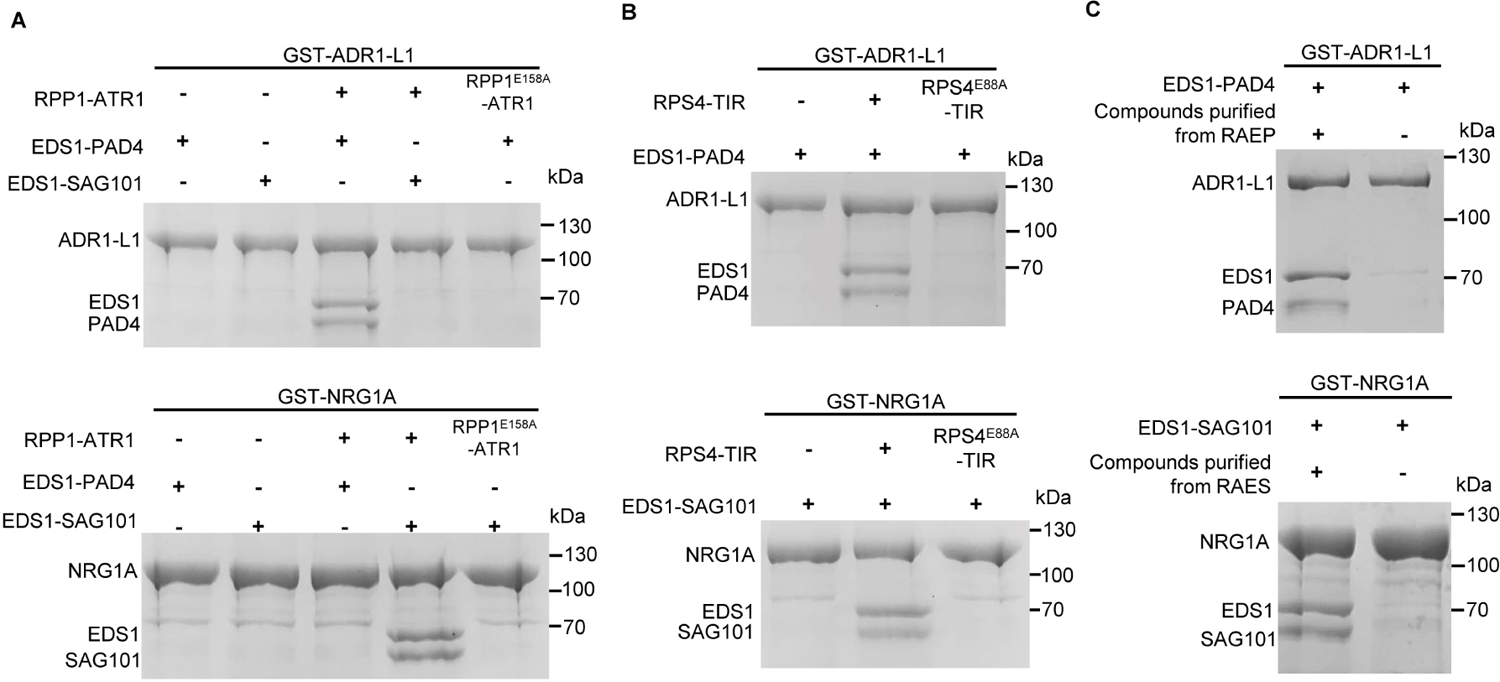
TIR-catalyzed products induce EDS1 heterodimer interaction with hNLRs in insect cells. (**A**) RPP1-ATR1 induces EDS1-PAD4 and EDS1-SAG101 interaction with ADR1-L1 and NRG1A, respectively, in insect cells. N-terminally GST-tagged ADR1-L1 or NRG1A was co-expressed with non-tagged EDS1, C-terminally strep-tagged PAD4 or SAG101, RPP1 (wild type or catalytic mutant E158A) and 10xHis-tagged ATR1 in insect cells. GST-ADR-L1 or NRG1A was purified using Glutathione Sepharose 4B (GS4B) beads. GS4B-bound proteins were eluted, separated by SDS-PAGE and detected by Coomassie Brilliant Blue staining. (**B**) RPS4 TIR (residues 1-236) induces EDS1-PAD4 and EDS1-SAG101 interaction with ADR1-L1 and NRG1A, respectively, in insect cells. Experiments were performed as described in (A) except that RPP1 and ATR1 were replaced with RPS4 TIR. (**C**) EDS1-PAD4 or EDS1-SAG101 was expressed as described in (A) and purified with Strep-Tactin resin. The purified protein was concentrated and denatured by heating. After centrifugation, the supernatant was collected and incubated with separately purified N-terminally GST-tagged ADR1-L1 or NRG1A and apo-EDS1-PAD4 or apo-EDS1-SAG101. The mixture was passed over GS4B beads and bound proteins were eluted, analyzed by SDS-PAGE and detected by Coomassie Brilliant Blue staining.

We next investigated whether products of TIR enzyme activity bind to and support EDS1-PAD4 and EDS1-SAG101 interactions with hNLRs *in vitro*. For this we co-expressed RAEP in insect cells, and the EDS1-PAD4 complex thus expressed is predicted to contain the products catalyzed by the RPP1 resistosome. We purified the EDS1-PAD4 protein and extracted the presumed small molecules by denaturing the complex protein. The extracted small molecules induced interaction of separately purified apo-EDS1-PAD4 with ADR1-L1 (Fig. 1C). Similarly, small molecules extracted from the RAES-purified and denatured EDS1-SAG101 stimulated interaction of separately purified apo-EDS1-SAG101 with NRG1A (Fig. 1C). These biochemical data show that enzymatic products of TIR proteins interact with and facilitate EDS1-PAD4 and EDS1-SAG101 specific interactions with hNLRs.

## Structures of TIR-catalyzed small molecules activating EDS1-PAD4

We next determined the identities of the extracted small molecules promoting EDS1-PAD4 interaction with ADR1-L1 using liquid chromatography coupled with high resolution mass spectrometry (LC-HRMS). Two singly charged ions with mass-to-charge ratios (m/z) 558.0657 (-) and 638.0331 (-) were observed in the compounds extracted from RAEP but not control samples (fig. S1A). To determine the chemical structures of these two compounds, we purified EDS1-PAD4 by co-expressing RAEP in insect cells, crystallized the EDS1-PAD4 complex and solved its structure. The crystal structure of the complex was solved with molecular replacement using EDS1-SAG101 (*15*) as the template, and was refined to 2.29 Å resolution (Fig. 2A and Table S1).

**Fig. 2.**
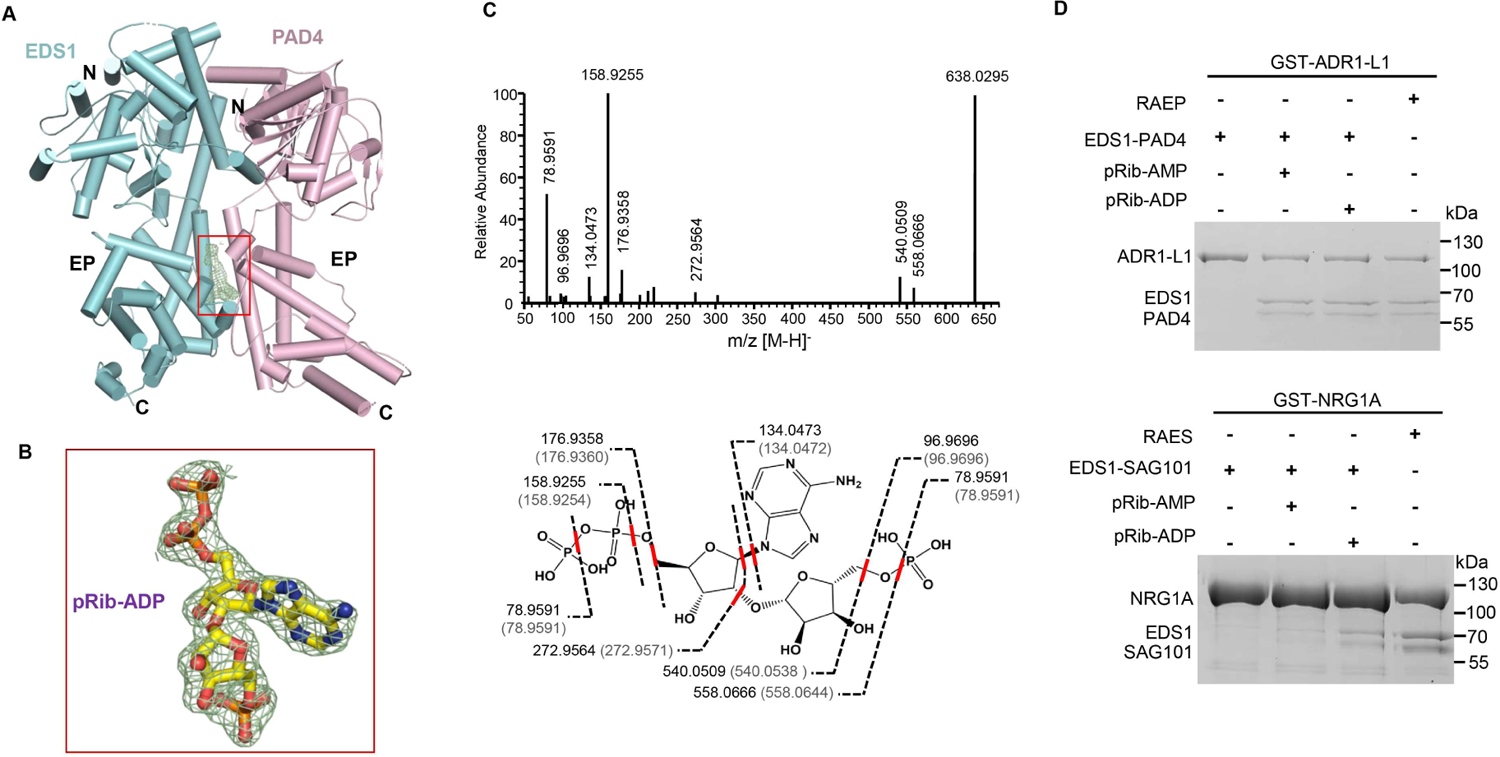
Structural determination of the EDS1-PAD4 complex with bound pRib-ADP. (**A**) Crystal structure of EDS1-PAD4 bound by RPP1-catalyzed small molecule. Highlighted within the red frame is omit electron density (*2Fo-Fc* contoured at 1.5 sigma) between the EP domains of EDS1 and PAD4. (**B**) Close-up view of the electron density highlighted in (**A**) with modeled pRib-ADP (in stick). (**C**) Mass analysis of small molecules bound to EDS1-PAD4. Top: MS/MS spectra after CID fragmentation of the ion with m/z = 638.0295 (z =1^-^). Bottom: Scheme showing the proposed routes of formation of various ions shown above. Numbers in parentheses indicate theoretical molecular weights of proposed ions. (**D**) Synthetic pRib-AMP and pRib-ADP induce strong EDS1-PAD4 and weak EDS1-SAG101 interaction with ADR1-L1 and NRG1A, respectively. GS4B beads bound by GST-ADR1-L1 or GST-NRG1A were incubated with excess purified apo-EDS1-PAD4 or apo-EDS1-SAG101 and synthetic pRib-AMP/ADP on ice for 30 min. After extensive washing, proteins bound to GS4B beads were eluted, analyzed by SDS-PAGE and detected by Coomassie Brilliant Blue staining.

In the final electron density map, there is a segment of unambiguous electron density between the partner EP domains which remains unoccupied after modeling of EDS1 and PAD4 (Fig. 2B). The ion feature with m/z 558.0657 (-) from LC-HRMS suggested adenosine diphosphate ribose (ADPR, molecular mass 559.0717, 2.4 ppm mass shift) within 3 ppm (parts per million) as one of the small molecules. However, ADPR did not match the unfilled electron density, suggesting that the small molecule bound by EDS1-PAD4 is an ADPR isomer or isomer derivative. This is consistent with LC-HRMS analysis showing that ADPR had a retention time different from that of the small molecule with m/z 558.0657 (-) (fig. S1B). One ADPR isomer, 2’-(5’’-phosphoribosyl)-5’-adenosine monophosphate (pRib-AMP), fitted the electron density well except where a small portion connecting the terminal phosphate group remains void (fig. S2C). The ion feature with m/z 638.0331 (-), with a mass shift of 79.967 from m/z 558.0657 (-), suggested a molecule with an additional phosphate group on pRib-AMP. Adding one phosphate group to the terminal void density produced 2’-(5’’-phosphoribosyl)-5’-adenosine diphosphate (pRib-ADP) which fitted into the electron density map very well (Fig. 2B). A higher pRib-ADP affinity for EDS1-PAD4, might explain why pRib-ADP was captured in the crystal structure of EDS1-PAD4.

To verify the assignment of pRib-AMP and pRib-ADP in the EDS1-PAD4 complex, standard compounds were chemically synthesized. MS/MS spectra and chromatographic elution profiles, confirmed the identification of pRib-AMP and pRib-ADP as EDS1-PAD4 ligands (Fig. 2C and fig. S2A, B and D). We tested whether synthetic pRib-AMP and pRib-ADP induce EDS1-PAD4 and EDS1-SAG101 interactions, respectively, with ADR1-L1 and NRG1A. Both compounds promoted EDS1-PAD4 interaction with ADR1-L1, similar to that detected after co-expression of ADR1-L1 with RAEP (Fig. 2D, top). The small molecules were much less efficient in inducing EDS1-SAG101 interaction with NRG1A (Fig. 2D, bottom). These structural and biochemical data demonstrate that pRib-AMP/ADP are TIR-catalyzed metabolites that preferentially induce EDS1-PAD4 interaction with ADR1-L1.

## Recognition mechanism of pRib-ADP by EDS1-PAD4

In the crystal structure, pRib-ADP adopts an extended conformation interacting with a pocket formed by the EDS1 and PAD4 EP domains (Fig. 3A). In support of the structural observation, the pRib-ADP binding pocket overlaps with positively charged EP-domain surfaces of EDS1 and PAD4 that are indispensable for EDS1 immunity signaling (*15, 16, 20*). pRib-ADP binding results in complete burial of the small molecule except its terminal phosphate group (Fig. 3A). An extensive network of hydrogen bonds, including those mediated by several well-ordered water molecules, appears to govern binding of pRib-ADP to the pocket (Fig. 3B), indicating specific recognition of pRib-ADP by EDS1-PAD4. The 5-phospho-beta-D-ribosyl moiety of pRib-ADP is recognized by EDS1 residues Asp469, Asn472 and Tyr473. The phosphate group of this moiety also forms salt bridges with Arg314 and Arg425 and van der Waals contact with Phe387 of PAD4 (Fig. 3B). Thus, both EDS1 and PAD4 make significant contributions to recognition of the ADP moiety of pRib-ADP. The adenine group stacks against PAD4 Tyr383 at one side and EDS1 Tyr473 at the other, and is further stabilized by hydrogen bonds with EDS1 Thr482. Besides making van der Waals contacts with PAD4 Lys379/Lys380, the ADP moiety ribose group hydrogen bonds with EDS1 Arg493 directly and via a water molecule. The two phosphate groups of the ADP moiety form in total eight hydrogen bonds with their neighboring residues - five with EDS1 and three with PAD4 (Fig. 3B, right). Structure-based sequence alignments indicated that EDS1 and PAD4 residues involved in recognition of pRib-ADP are highly conserved in different plant species (fig. S3), suggesting that pRib-AMP/ADP binding is an evolutionarily conserved activity of EDS1-PAD4. A cyclic ADPR is less likely to fit in the pRib-ADP-binding pocket in EDS1-PAD4, although the precise chemical structure of v-cADPR (*12*) remains undefined. In *Arabidopsis*, mutations in EP-domain surface residues EDS1 Arg493, Lys478 and PAD4 Arg314, Lys380 identified here as contacts for pRib-ADP binding, did not disrupt heterodimer formation but abolished TNL effector-triggered and basal immunity (*16, 18, 20*). Also, mutation of PAD4 Arg314 disabled TNL-induced EDS1-PAD4 association with ADR1 in *Arabidopsis* (*20*).

**Fig. 3.**
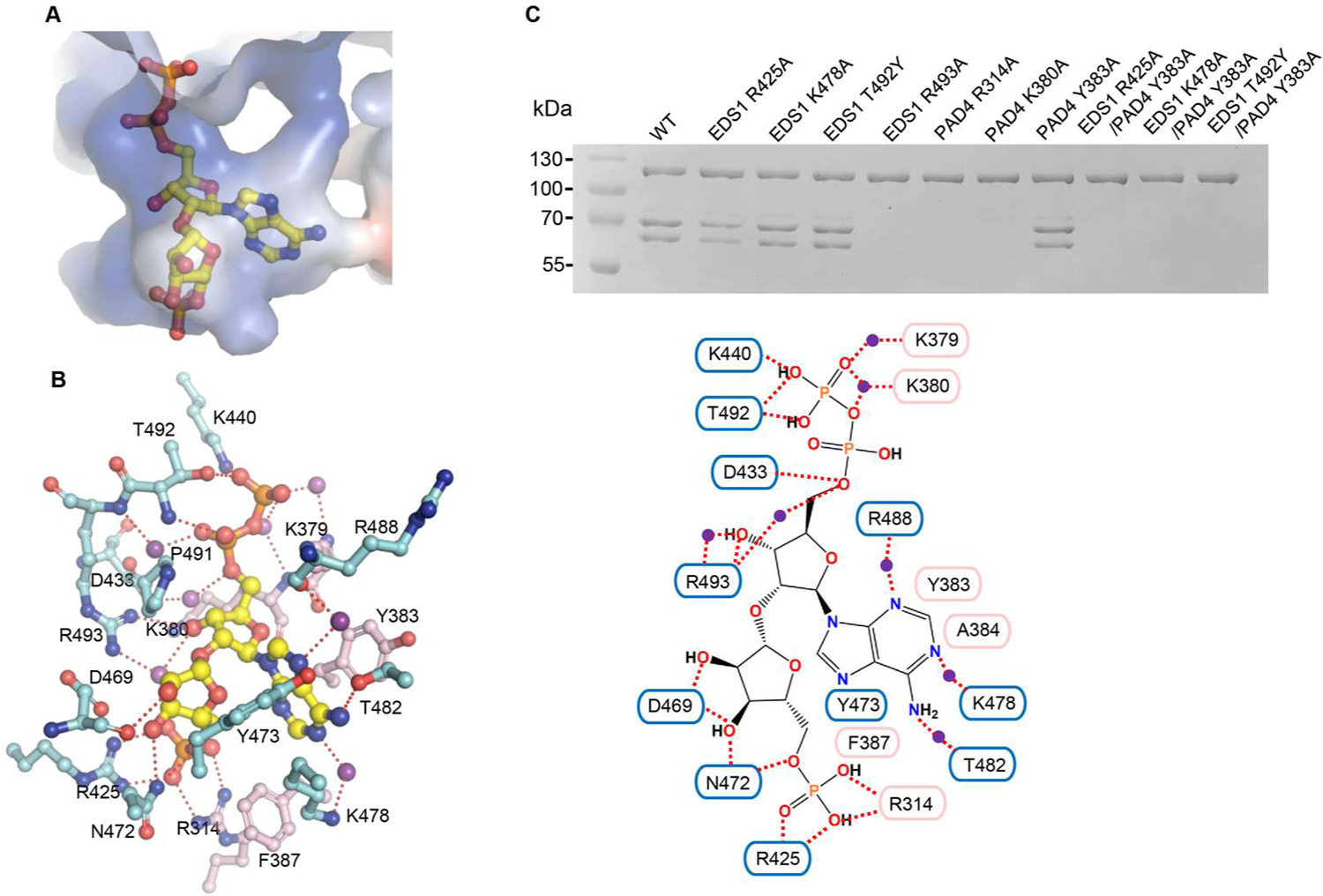
Specific recognition of pRib-ADP by EDS1-PAD4. (**A**) Close-up view of the pRib-ADP-binding pocket of EDS1-PAD4. EDS1-PAD4 is shown in electrostatic surface and pRib-ADP in stick. (**B**) Left: EDS1-PAD4 EP-domain amino acid coordinates for pRib-ADP. Dashed lines represent polar interactions. Purple spheres represent water molecules. Right: Diagram showing interaction between EDS1-PAD4 and pRib-ADP. Cyan and purple frames represent residues from EDS1 and PAD4, respectively. (**C**) Mutagenesis analysis of the EDS1-PAD4 pRib-ADP binding pocket. Assays were performed as described in Fig. 1A.

To verify the pRib-ADP interaction with EDS1-PAD4 observed in the crystal structure, we engineered amino acid substitutions at the pRib-ADP binding pocket in EDS1 and PAD4 and tested their effects on EDS1-PAD4 interaction with ADR1 induced by the RPP1 resistosome in insect cells. Mutations of positively charged residues EDS1 Arg493, PAD4 Arg314 and PAD4 Lys380 coordinating the phosphate group of the pRib moiety had little effect on EDS1-PAD4 dimer formation but markedly reduced EDS1-PAD4 interaction with ADR1-L1 (Fig. 3C). When combined with EDS1 R425A, EDS1 T492Y or PAD4 Y383A, mutation of PAD4 Tyr383 stacking the adenosine moiety resulted in a complete loss of the interaction.

## Activation of TIR-containing proteins promotes pRib-ADP/AMP accumulation in plants

We investigated whether activation of TIR-containing proteins leads to pRib-ADP/AMP production in plants. *Arabidopsis* TIR-only protein RBA1 (*30*) was used because it promotes EDS1-PAD4 interaction with ADR1-L1 in an *Nb* tobacco transient co-expression assay (fig. S4) (*26*). LC/MS detection of these two compounds upon RBA1 expression in a signaling defective *Nb* tobacco *epss* line lacking all EDS1-family proteins (*Nb-epss*) (*16*) did not produce clear signals, possibly due to pRib-ADP/AMP instability in leaves. Indeed, LC/MS assays showed that ∼50% of pRib-AMP and more than 90% pRib-ADP was degraded within one hour of mixing with *Nb epss* tissue lysate (fig. S5). Since binding to EDS1-PAD4 might protect pRib-AMP/ADP from hydrolysis (Fig. 2A), *Nb*-*epss* leaves expressing RBA1 were co-sonicated with insect cells expressing EDS1 and PAD4 to detect EDS1-PAD4-captured pRib-AMP/ADP with LC/MS. Plants expressing wild type RBA1 produced strong pRib-AMP signals in the assay (Fig. 4 and fig. S6). By contrast, expression of an RBA1 NADase catalytic mutant (E86A) substantially reduced pRib-AMP accumulation (Fig. 4 and fig. S6). We failed to observe accumulation of pRib-ADP under the same conditions. This unstable small molecule might need to bind to EDS1-PAD4 immediately upon generation to evade degradation in plants. These data show that pRib-AMP produced in TIR-activated plant tissues binds to the EDS1-PAD4 complex.

**Fig. 4.**
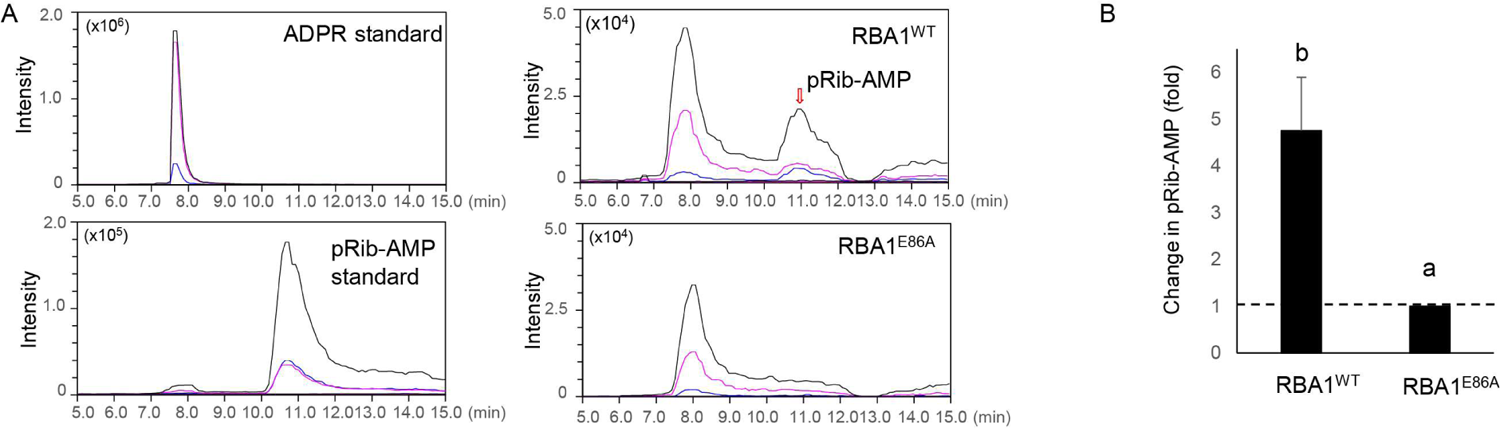
Expression of TIR-only protein RBA1 induces pRib-AMP accumulation in *Nb* tobacco. (**A**) MRM analyses of pRib-AMP in *Nb* tobacco. *Nb epss* leaves expressing RBA1 or catalytic mutant *RBA1*^E86A^ and insect cells expressing apo-EDS1-PAD4 complex were mixed and co-sonicated. The apo-EDS1-PAD4 complex in the lysate was purified through affinity chromatography and LC/MS was used to analyze small molecules bound in the purified apo-EDS1-PAD4 complex. The left two panels are MRM chromatograms for ADPR and synthetic pRib-AMP, respectively. The right two panels are representative chromatograms of pRib-AMP captured by EDS1-PAD4 in *Nb epss* expressing wild type *RBA1* and*RBA1*^E86A^, respectively. Black, purple and blue lines respectively indicate different transitions, 560.10>136.20, 560.10>348.00 and 560.10>97.20. (**B**) Relative change in pRib-AMP accumulation in *Nb epss* plants expressing RBA1 or RBA1^E86A^. The statistics were based on LC-MS results shown in and fig. S6. The amount of pRib-AMP in RBA1^E86A^-expressing *Nb epss* was normalized to 1.0.Significance was calculated with Tukey′s HSD test (n = 3, α= 0.05; different letters indicate a significant difference).

## Allosteric activation of EDS1-PAD4 by pRib-ADP/pRib-AMP

Having established that pRib-AMP/ADP are ligands of EDS1-PAD4, we examined the receptor complex activation mechanism by these two compounds. For this, we purified the apo-EDS1-PAD4 heterodimer from insect cells and solved a cryo-EM structure of the complex at 2.93 Å resolution (Fig. 5A, fig. S7 and Table S2). The EDS1 structure is nearly identical to that in the crystal structure of pRib-ADP bound EDS1-PAD4 (fig. S8A). As predicted, the structures of the PAD4 N-terminal lipase-like and C-terminal EP domains largely resemble their SAG101 counterparts (fig. S8B), although relative positioning of the two structural domains is strikingly different (fig. S8C). Structure alignment between the free and pRib-ADP-bound forms of EDS1-PAD4 reveals that PAD4 undergoes a marked conformation change in its C-terminal EP domain upon pRib-ADP binding (Fig. 5, B and C). Compared to the PAD4 apo-form, the PAD4 EP domain in its ligand-bound state is rotated ∼10 degrees anti-clockwise (viewed from the bottom) (Fig. 5C). This structural change leads to dissolving of the central part of a long helix connecting the PAD4 lipase-like and EP domains, which in turn allows movement of the C-terminal half of the helix towards EDS1 (Fig. 5, A and B). Similar conformation changes occur at the PAD4 short α-helix (α15) and N-terminal portion of α17. Collectively, these changes create a pRib-ADP-binding site between the EP domains of PAD4 and EDS1 (Fig. 5B), and suggest that an induced-fit mechanism underlies small molecule binding. Except for the terminal phosphate group, pRib-ADP becomes solvent inaccessible following binding to EDS1-PAD4 (Fig. 3A). Therefore, the small molecule is unlikely to be directly involved in interaction with ADR1-L1 but rather mediates conformational changes in the PAD4 EP domain enabling ADR1 binding to EDS1-PAD4. We conclude that pRib-AMP or pRib-ADP binding allosterically activates the EDS1-PAD4 receptor complex to interact with ADR1-family hNLRs. Based on our data, we propose a model in which TIR-generated pRib-AMP/ADP induces activation of EDS1-PAD4 to allosterically promote its association with ADR1 and plant immunity (fig. S9A).

**Fig. 5.**
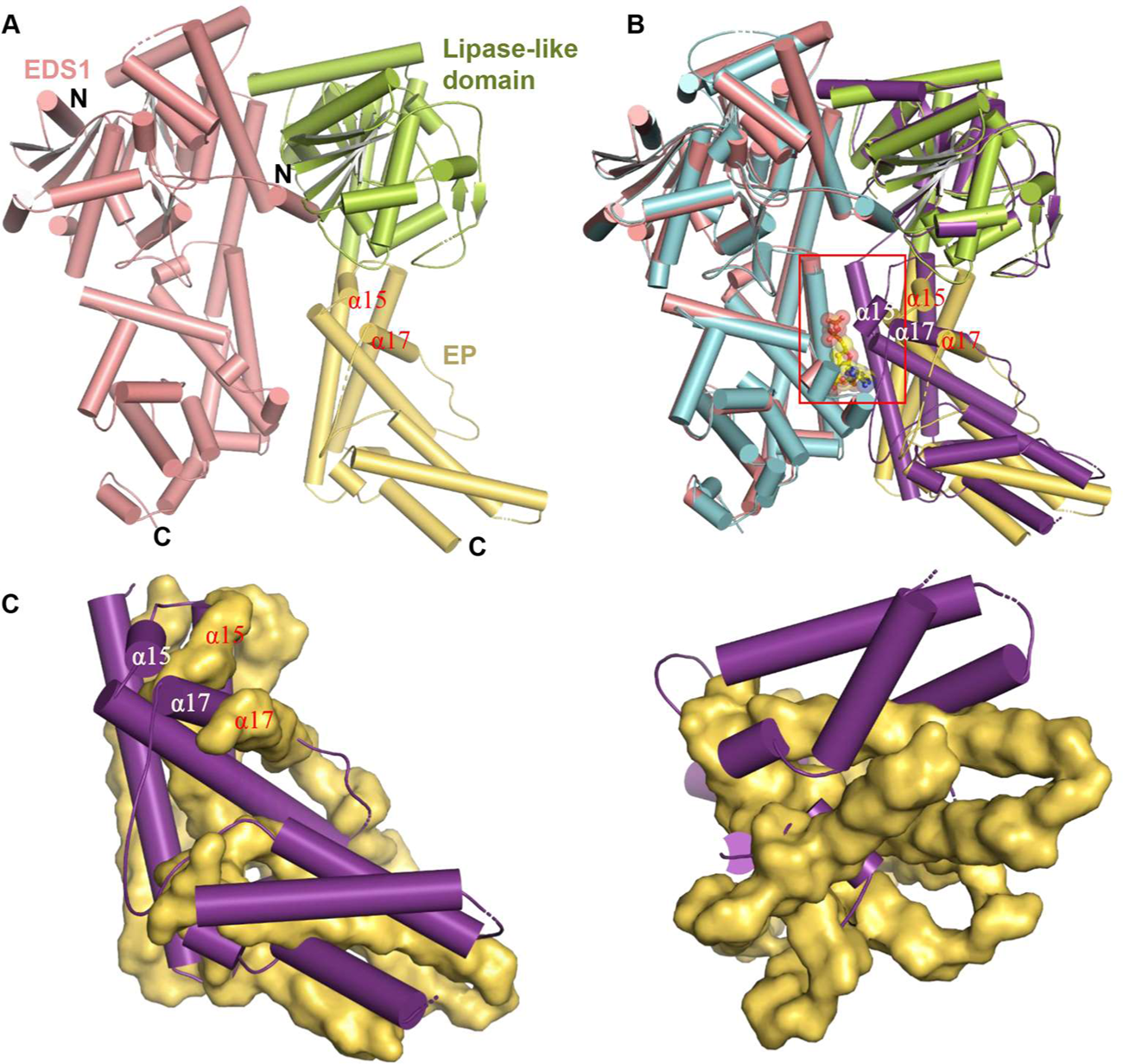
Allosteric activation of EDS1-PAD4 by pRib-ADP/AMP. (**A**) Cryo-EM structure of apo-EDS1-PAD4. PAD4 lipase-like and EP domains are shown in green and light orange, respectively. (**B**) Structural alignment between apo- and pRib-ADP-bound forms of EDS1-PAD4. Highlighted with an open red frame is the pRib-ADP-binding pocket between the EDS1 and PAD4 EP domains. EDS1 and PAD4 in the ligand-bound form are shown in cyan and purple, respectively. EDS1 in apo-EDS1-PAD4 was used as the template for structural alignment. Color codes for the apo-EDS1-PAD4 are the same as those in (A). The secondary structure elements labelled in white and red are from ligand-bound and apo-EDS1-PAD4, respectively. (**C**) Two orientations showing structural alignment of the EP domains of PAD4 in apo-(in surface) and ligand-bound (in cartoon) forms of EDS1-PAD4. Left: the orientation is the same as that shown in (B). Right: bottom view of the structural alignment.

## Discussion

Plant TIR NADase-generated small molecule signaling intermediates proposed to activate immune responses have been elusive. One challenge has been to develop an *in vitro* biochemical assay that detects TIR-catalyzed bioactive molecules. Here, we reconstituted EDS1 heterodimer associations with hNLRs induced by different TIR-containing proteins in insect cells (Fig. 1, A and B). Expression of TIR-only protein RBA1 in *Nb* tobacco promoted accumulation of pRib-AMP in a catalytic activity-dependent manner (Fig. 4, A and B). These results highlight pRib-AMP/ADP as potentially conserved small molecules for EDS1 signaling in seed plants. This is supported by the high degree of conservation of pRib-ADP coordinating residues in EDS1 and PAD4 (fig. S3) and triggering of *EDS1*-dependent cell death in *Nb* tobacco plants by a monocot *TIR*-only protein (*12*). The terminal phosphate group of pRib-ADP is solvent-exposed when bound to EDS1-PAD4 (Fig. 3A). It is therefore possible that pRib-ADP derivatives with covalent modifications to this phosphate group are also recognized by EDS1-PAD4. Biochemical and structural data reveal that *Arabidopsis* EDS1-PAD4 functions as a receptor complex for pRib-AMP/ADP (Figs 1 and 3). Supporting the biological significance of pRib-AMP/ADP recognition by EDS1-PAD4, mutations disrupting pRib-AMP/ADP binding (Fig. 3C) suppress EDS1-PAD4-mediated pathogen resistance (*15, 18, 20*). pRib-AMP/ADP binding induces conformational changes in EDS1-PAD4 to allosterically promote its association with ADR1-L1 (Fig. 5). The above characteristics lead us to propose that TIR-catalyzed products pRib-AMP/ADP act as second messengers in plant immunity signaling. The mode of action of these two compounds is reminiscent of cyclic nucleotide-based second messengers such as cyclic GMP-AMP (cGAMP) in animals (*31*) (fig. S9B).

Besides mediating TNL receptor signaling, the *Arabidopsis* EDS1-PAD4 dimer promotes defense reprogramming downstream of a non-TNL-type intracellular NLR RPS2 (*18, 32*) and cell-surface immune receptor RLP23 (*14, 33, 34*), via what we identify here as the pRib-AMP/ADP binding pocket. Thus, pRib-AMP/ADP likely have a broad role in biotic stress signaling, consistent with the molecules being second messengers (*12, 34*). In further comparison to well-established second messengers such as phosphatidylinositol 3,4,5-trisphosphateinositol (PIP_3_) and cADPR (*35*), pRib-AMP/ADP convert externally generated stimuli (pathogen effectors) to intracellular Ca^2+^ signals, potentially by activating the Ca^2+^-permeable channel of ADR1 (*27*). Second messenger signaling is generally terminated by metabolic enzymes (*36*). Although the processes that remove pRib-ADP/AMP are not known, these molecules are rapidly degraded in *Nb* tobacco extracts (fig. S5).

The current data show that TIR-containing proteins catalyze production of pRib-AMP/ADP which directly activate EDS1-PAD4 dimers. TIR-containing proteins can also encode a 2’,3’-cAMP/cGMP synthetase activity promoting *EDS1*-dependent cell death. Thus, plant TIR-containing proteins appear to constitute a large family of enzymes producing noncanonical and nucleotide-based signaling molecules for regulation of immune and possibly other stress responses. We previously proposed that 2’,3’-cAMP/cGMP facilitate self-amplification of EDS1 signaling (*37*). An outstanding question is how TIR-generated pRib-AMP/ADP or their derivatives intersect with 2’,3’-cAMP/cGMP in driving the plant immune response.

## Acknowledgments

We thank the National Protein Science Facility at Tsinghua University for technical assistance. We thank the staff of Beamline BL02U1, BL19U at SSRF for assistance in data collection. We thank J. Lei, X. Li, X. Fan, and N. Liu at Tsinghua University for data collection. We acknowledge the Tsinghua University Branch of the China National Center for Protein Sciences (Beijing) for providing the cryo-EM facility support and the computational facility support on the cluster of Bio-Computing Platform. A patent application (CN 202210087257.X) has been filed relating to this work.

## Supplementary Materials

### Materials and methods

#### Protein expression and purification

For purification of the apo-EDS1-PAD4, constructs of full-length *Arabidopsis* EDS1 (residues 1-623, cloned in a pFastbac 1 vector) and full length PAD4 (residues 1-541, cloned in a modified pFastbac 1 vector with a C-terminal twin-StrepII tag) were co-expressed in Sf21 at 28°C, using the Bac-to-Bac baculovirus expression system (Invitrogen) protocols. After 48 h of recombinant baculovirus infection, cells were harvested and re-suspended in buffer containing 25 mM Tris (pH 8.0), 150 mM NaCl and 1 mM PMSF. The harvested cells were lysed by sonication, the lysate centrifuged and the supernatant collected. The apo-EDS1-PAD4 complex protein was purified using Streptactin Beads (Novagen) from the supernatant. Bound proteins were eluted in a buffer containing 25 mM Tris (pH 8.0), 150 mM NaCl, 2.5 mM desthiobiotin and further purified by size exclusion chromatography (Superdex 200 increase 10/300, GE Healthcare) in a buffer containing 25 mM Tris (pH 8.0), 150 mM NaCl.

To purify EDS1-PAD4 bound by TIR-generated small molecules, the EDS1 and PAD4 constructs described above were co-expressed with RPP1_WsB (residues 61-1221, cloned in a modified pFastbac 1 vector with a C-terminal twin-StrepII tag) and ATR1_Emoy2 (residues 52-311, cloned in a modified pFastbac 1 vector with a C-terminal 10x His tag) in Sf21 at 28°C. A similar procedure was used to purify the EDS1-PAD4 complex protein.

#### Reconstitution of EDS1-PAD4-ADR1-L1/EDS1-SAG101-NRG1A interaction in insect cells

To reconstitute TIR-activated ADR1, N-terminally GST-tagged ADR1-L1 (residues 30-816, cloned in a pFastbac 1 vector with a N-terminal GST tag) was co-expressed with non-tagged EDS1, C-terminally strep-tagged PAD4 with RPP1 (residues 61-1221) and 10xHis-tagged ATR1 in Sf21 at 28 °C using Bac-to-Bac baculovirus expression system (Invitrogen) protocols. Insect cells were harvested, lysed and centrifuged. GST-ADR1-L1 protein was purified using Glutathione Sepharose 4B beads (∼200 µl, Novagen). The resin was washed three times with 1.0 ml buffer containing 25 mM Tris (pH 8.0), 150 mM NaCl. Bound proteins were eluted in buffer containing 25 mM Tris (pH 8.0), 150 mM NaCl and 15 mM GSH. The eluted proteins were separated by SDS-PAGE and detected by Coomassie Brilliant Blue staining. A similar procedure was used to reconstitute EDS1-SAG101-NRG1A (residues 58-810, cloned in a pFastbac 1 vector with a N-terminal GST tag) interaction. EDS1-PAD4-ADR1-L1 or EDS1-SAG101-NRG1A interaction was also reconstituted using non-tagged RPS4 TIR domain (residues 1-236) in place of RPP1 and ATR1, using similar protocols.

#### Pull-down assay

EDS1-PAD4 protein bound to RPP1 resistosome-catalyzed small molecules was purified and concentrated to 10.0 mg/ml. The concentrated protein (400 µl) in a 1.5-ml centrifuge tube was denatured by heating (92 °C for 7 min). The sample was centrifuged to remove precipitate and the supernatant collected to repeat the heating-centrifuge steps two more times. The final 400 µl supernatant was divided in two in 1.5-ml centrifuge tubes. N-terminally GST-tagged ADR1-L1 and N-terminally GST-tagged NRG1A were expressed in Sf21 insect cells and purified using GS4B beads as described above. The GS4B beads bound by GST-tagged ADR1-L1 were incubated with excess separately purified apo-EDS1-PAD4 and 200 µl supernatant on ice for 30 min. The GS4B beads were then washed with 1.0 ml buffer containing 25 mM Tris (pH 8.0), 150 mM NaCl 3 times. Bound proteins were eluted in buffer containing 25 mM Tris (pH 8.0), 150 mM NaCl and 15 mM GSH, analyzed by SDS-PAGE and detected by Coomassie Brilliant Blue staining. Similar protocols were used to assay GST-tagged NRG1A interaction with EDS1-SAG101.

The pulldown assay described above was used to test, respectively, EDS1-PAD4 and EDS1-SAG101 interaction with ADR1-L1 and NRG1A induced by chemically synthesized pRib-ADP/AMP. 200 µl of pRib-AMP (final concentration 0.5 mM) and 500 uM of pRib-ADP (final concentration 0.5 mM) were used in the assays.

#### Chemical synthesis of pRib-AMP/ADP

The synthesis of compound **15** started from *N*-benzoyladenosine (Data S1, Scheme 1). Selective silylation of nucleoside **1** followed by the reaction with beta-D-ribofuranose 1-acetate 2,3,5-tribenzoate gave intermediate **3**. After desilylation, the 3’-OH in **4** was selectively protected with a TBS group. The first phosphate group was introduced at the 5’-OH position using dibenzyl diisopropylphosphoramidite. Similarly, the second phosphate group was installed at the 5’’-OH position *via* five steps. Sequential desilylation and debenzylation of **13** produced compound **15**.

Selective protection of the 5’’-OH in **9** followed by benzoylation resulted in intermediate **16** (Data S1, Scheme 2). Then the protecting groups at the 5’’-OH position and at the phosphate moiety were removed sequentially and a phosphate group was incorporated into the above two sites, respectively. Removal of all the protecting groups in **19** *via* three steps produced compound **22**. All compounds were confirmed by NMR spectra (Data S1, NMR spectra of compounds).

#### LC-HRMS assay

Approximately 0.3 mg of purified apo-EDS1-PAD4 and small molecule-bound forms of EDS1-PAD4 were concentrated to 20 µl in 0.1M NH_4_HCO_3_ buffer (PH 7.6). Then, 80 µl methanol was added to the NH_4_HCO_3_ solution. The mixed solution was stored in a −80°C fridge for 30 min and then centrifuged to remove the precipitate. The supernatant was transferred to a new 1.5-ml centrifuge tube for LC-MS. Analysis was performed using an orbitrap mass spectrometer (Q-Exactive HFX, Thermo Fisher, CA) coupled with a Vanquish UHPLC system (Thermo Fisher, CA) in negative ion mode. Volume of 5 µL supernatant was loaded to a BEH Amide column (100 × 2.1 mm, Waters, USA) for LC separation. Mobile phase A was 95% acetonitrile with 5 mM ammonium acetate and mobile phase B was aqueous containing 5 mM ammonium acetate. The sample was then eluted by mobile phase B from 10% to 50% within 5 min. Mass range of m/z 400-700 in negative ion mode was used for data acquisition. Data dependent acquisition (DDA) was applied for MS/MS. Mass resolutions of full scan and MS/MS spectra were set as 70,000 and 17,500, respectively. The source parameters were as follows: spray voltage: 3,000v; capillary temperature: 320 °C; heater temperature: 300 °C; sheath gas flow rate: 35 arb; auxiliary gas flow rate: 10 arb. Data analysis was performed by the software Xcalibur 4.4 (Thermo Fisher, CA).

#### Crystallization, data collection, structure determination and refinement

Small molecule-bound EDS1-PAD4 complex protein was concentrated to about 7.0 mg/ml for crystallization. Crystallization of the complex was performed using hanging-drop vapor-diffusion by mixing 1 μl protein with 1 μl reservoir solution at 18°C. Diffraction quality crystals of the ligand-bound EDS1-PAD4 complex were obtained in the buffer containing 0.04 M citric acid, 0.06 M Bis-Tris propane pH 6.4, 20% w/v polyethylene glycol 3,350. All diffraction datasets were collected at Shanghai Synchrotron Radiation Facility (SSRF) on beam line BL19U using a CCD detector. Data were processed using the HKL2000 software package (*38*). The crystal structure of active EDS1-PAD4 complex was determined by molecular replacement (MR) with PHASER (*39*) using the structure of EDS1-SAG101 (PDB code: 4NFU) as the initial searching model. The model from MR was built with the COOT (*40*) and subsequently subjected to refinement by Phenix_real_space_refine. Statistics of diffraction data and refinement of the EDS1-PAD4 model are summarized in Table S1. Structural figures were prepared using PyMOL (http://www.pymol.org/).

#### Cryo-EM sample preparation and data acquisition

After gel filtration, the apo-EDS1-PAD4 complex was concentrated to ∼0.8-1 mg/mL for cryo-EM sample preparation. Quantifoil Au R 1.2/1.3 300 mesh holey carbon girds were glow-discharged for 30 s at medium level in Harrick Plasma after 2 min evacuation. A volume of 3 µL of protein solution was applied to the grids, and samples were vitrified in a FEI Vitrobot Mark IV (Thermo Fisher Scientific) using a blotting time of 2.5-3.5 s at 8 °C with humidity 100%.

Cryo-EM data were collected in counted super-resolution mode on a Titan Krios3 (FEI) electron microscope operating at 300 kV equipped with VPP (Volta Phase Plate), a Gatan Quantum energy filter and Gatan K3 direct detection camera at ×105,000 magnification, with a physical pixel size of 0.3373 Å. The slit width of the energy filter was set to 20 eV. Using AutoEMation data collection software in super-resolution mode (*41*), the micrographs were dose-fractionated into 32 frames with a total exposure time of 8 s and a total electron exposure of ∼ 50 electrons per Å^2^, with defocus values ranging from −1.3 to −1.8 µM.

#### Image processing and 3D reconstruction

The stacks were 2 × Fourier binned, dose-weighted and summed using MotionCor2 (*42*), resulting in a pixel size of 0.6746 Å per pixel. The contrast transfer function (CTF) was corrected using the CTFFIND4 and bad micrographs were manually removed (*43*). Further processing was performed using RELION-3.1 (*44–46*). The cryo-EM processing workflow for apo-EDS1-PAD4 is outlined in fig. S2. In total 3,695 stacks were collected and 1,964,771 particles were auto-picked using Laplacian-of-Gaussian in RELION-3.1. Several rounds of 2D classification were performed to remove junk particles. After 2D classification, good particles were used to generate an ab initio map in RELION-3.1. The initial model was used as a reference for a first round of 3D classification. 551,861 good particles were then selected for a second round of 3D classification. The most homogeneous particles were selected for final 3D auto-refinement. After CTF refinement and Post-Processing, the final resolution of the 3D reconstruction was 2.93 Å, based on gold standard FSC 0.143 criteria (*47*). Local resolution distribution was evaluated using RELION (*48*).

#### Model building and refinement

The crystal structure of pRib-ADP-bound EDS1-PAD4 was docked into the EM density in Chimera to build the model (*49*). Local differences were manually built or adjusted in Coot (*50*). Models were refined using multiple rounds of Phenix_real_space_refine (*51*). Model validation was performed with MolProbity and EMRinger (*51*). Model statistics are summarized in Table S2.

### Protein expression, co-immunoprecipitation and immunoblotting analyses

Overexpression of *Arabidopsis* TIR-only protein RBA1 was used to induce association of EDS1-PAD4 with ADR1-L1 in *Nb* tobacco quadruple mutant *eds1a pad4 sag101a sag101b* (*Nb epss*). Agrobacteria were induced for two hr in Agromix (10 mM MgCl_2_, 10 mM MES pH 5.6, 150 μM acetosyringone) and syringe-infiltrated at OD_600_ = 0.2, or OD_600_ = 0.4 for RBA1, into *Nb epss* leaves. After 48 h, five 10 mm leaf discs were collected from each sample and protein extraction and co-immunoprecipitation was performed as described before (*19*). In short, total protein was extracted from ground plant material by addition of 2 mL extraction buffer (10% glycerol, 100 mM Tris–HCl pH 7.5, 5 mM MgCl_2_, 300 mM NaCl, 10 mM DTT, 0.5% IGEPAL CA-630, 2% PVPP, 1x Plant protease cocktail (11873580001, MilliporeSigma)), incubation for 10 min rotating and centrifugation for 35 min at 4500 x *g*. Extracts were filtered through two layers Miracloth (475855, MilliporeSigma) and 50 µL aliquots of this filtered extract were taken as input samples. Co-IPs were conducted for 2 h with 20 µL α-GFP trap agarose beads (gta-20, Chromotek) under constant rotation. Beads were collected by centrifugation at 800 x *g* for 5 min and washed 3 times in extraction buffer (without DTT and PVPP). All co-IP steps were performed at 4 °C. Beads and input samples were boiled at 95 °C in 2 × Laemmli buffer (100 µL were added to beads) for 10 min. Antibodies used for immunoblotting were α-GFP (11814460001, MilliporeSigma), α-FLAG (f1804, MilliporeSigma) and HRP-conjugated α-mouse antibody (A9044, Sigma-Aldrich). All antibodies were used in dilution 1:5000 (TBST with 3% non-fat milk powder).

### Detection of TIR-catalyzed pRib-AMP/ADP in *Nb* tobacco

RBA1 (residues 1-363) was cloned into pENTR/D-TOPO vector (Thermo Fisher Scientific, K240020). ThepENTR/D-TOPO *RBA1*^WT^ and *RBA1*^E86A^ plasmids were LR-recombined into pXCSG vectors with a C-terminal Myc tag. The generated constructs were transformed into *Agrobacterium* GV3101. *RBA1*^WT^ and *RBA1*^E86A^ were transiently expressed in *Nb* tobacco *epss* leaves by agroinfiltration. Final OD600 values for *RBA1*^WT^ and *RBA1*^E86A^ strains were set to 0.8. An RNAi silencing suppressor P19 was co-infiltrated with OD600 of 0.2. *Nb* tobacco *epss* leaves were harvested 48 h after infiltration, snap frozen in liquid nitrogen and ground to fine powder.

*Nb* tobacco *epss* leaf powders of *RBA*^WT^ or *RBA*^E86A^ were mixed with insect cells of EDS1-PAD4 complex in buffer containing 50 mM Tris-HCl pH 8.0, 150 mM NaCl (50 g *RBA*^WT^- or *RBA*^E86A^-expressing *Nb* tobacco *epss* leaves mixed with 1.5 L insect cells). The mixture of leaf powder and insect cells was lysed by sonication. Lysates were centrifuged at 30,000 *g* for 150 min. The supernatants were collected and allowed to flow through Ni-NTA resin. The Ni-NTA resin was first washed with 20 CV of wash buffer (50 mM Tris-HCl pH 8.0, 150 mM NaCl, and 15 mM imidazole) followed by 10 CV of wash buffer (100 mM NH_4_HCO_3_). Proteins bound in the resin were eluted with 2 CV Elute buffer (100 mM NH_4_HCO_3_ and 200 mM imidazole). Eluted EDS1-PAD4 complex protein was concentrated using 30K Centrifugal Filter (Merck Millipore) to 20 mg/ml. The buffer was changed to 100 mM NH_4_HCO_3_ during concentration. The purified EDS1-PAD4 complex protein was denatured with methanol at −80 °C for 1 h and then centrifuged to remove precipitate. The supernatant was used for LC-MS analysis.

Chromatography was performed on a Nexera XR 40 series HPLC (Shimadzu) using a Premier BEH zHILIC, 150×2.1 mm column (Waters). The column temperature was maintained at 50 °C and the sample tray at 4 °C. Samples (10 μl) were injected at a flow rate of 0.3 ml/min using 15 mM ammonium bicarbonate (Sigma) at pH 9.0, and 15 mM ammonium bicarbonate at pH 9.0, 90% acetonitrile (Fisher) as mobile phases A and B, respectively. Metabolites were eluted using the profile 0-8.5 min, 90-60% B; 8.5-9 min, 60-20% B; 9-16 min 20% B; 16-16.1 min, 5% B; 16.1-22 min, 5% B, 22-30 min, 90% B. The LC MS-8060 triple quadrupole mass spectrometer (MS) with electro spray ionization (Shimadzu) was operated in positive mode. Scheduled multiple reaction monitoring (MRM) was used to monitor analyte parent ion to product ion formation. MRM conditions were optimized using standard chemicals including: 559 ([M+H]^+^ 560.10>348.00, 560.10>136.25, 560.10>97.20), 639 ([M+H]^+^ 640.10>428.05, 640.10>136.15, 640.10>97.00). Both Q1 and Q3 quadrupoles were maintained in unit resolution. LabSolutions LCMS v5.97 software was used for data acquisition and LabSolutions Post run for processing (both Shimadzu). Metabolites were quantified by scheduled MRM peak intensity.

**Fig. S1.**
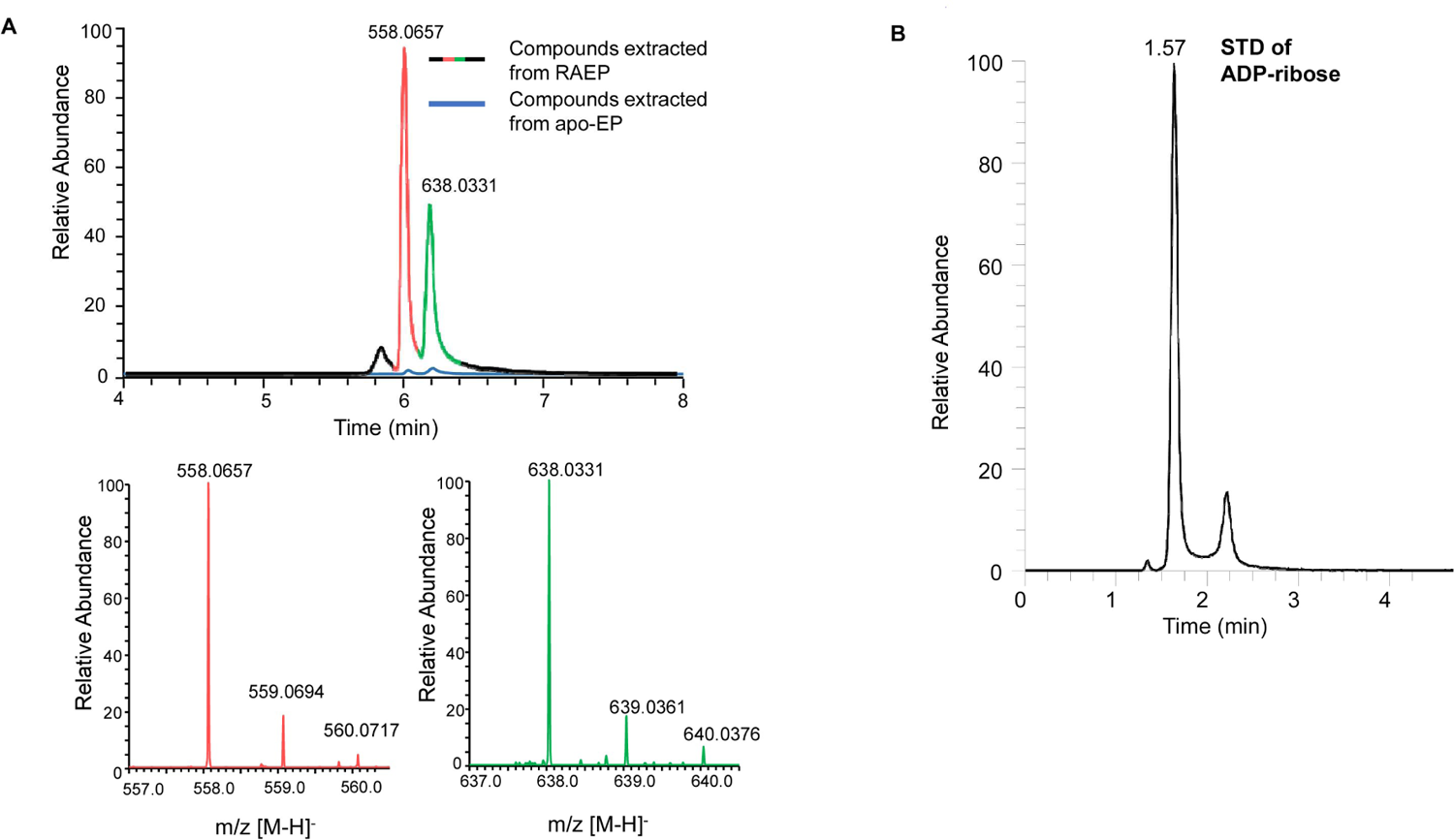
High resolution mass spectrometry (LC-HRMS) analysis of compounds bound to EDS1-PAD4. **(A)** Top: Chromatograms of supernatant extracts from the denatured apo-EDS1-PAD4 (blue) and EDS1-PAD4 co-expressed with RPP1-ATR1 (in colors). The purified proteins were denatured by 80% methanol(v/v) and after centrifuge, the supernatant was subjected to LC-HRMS analyses. The two peaks of LC from the supernatant of EDS1-PAD4 co-expressed with RPP1-ATR1 were shown in color with the red one corresponding to the ion mass of 558.0657 (-) and the green one to the ion mass of 638.0331 (-). Bottom two panels are isotopic distributions of the ions with m/z 558.0657 (-) and 638.0331 (-), respectively. (**B**) Chromatogram of ADP-ribose (ADPR) chemical standard from LC-HRMS. The retention time of ADPR was 1.57 min.

**Fig. S2.**
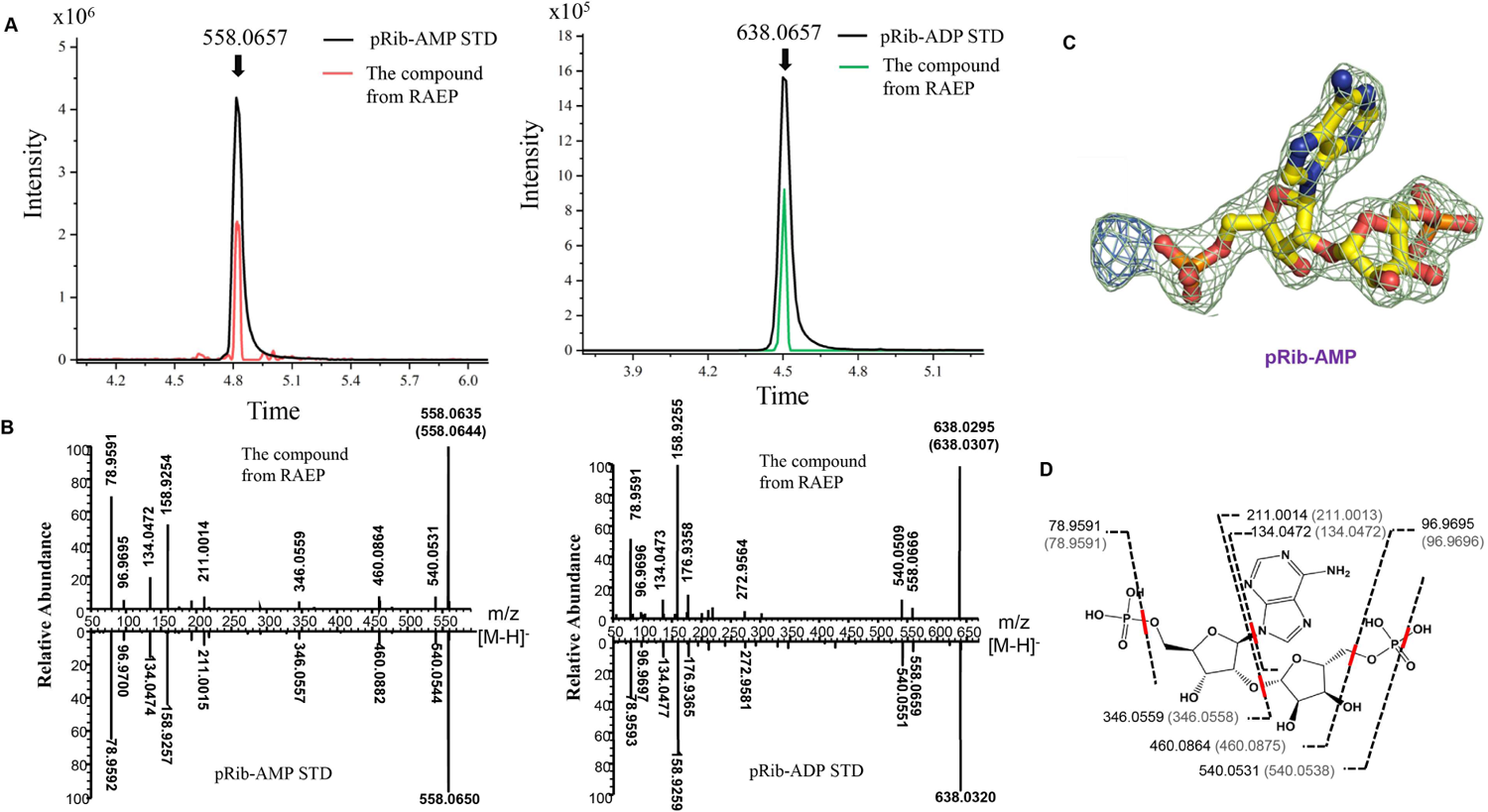
Confirmation of the identities of small molecules bound to EDS1-PAD4. **(A)** Left: chromatograms of the compound from EDS1-PAD4 co-expressed with RPP1-ATR1 (RPP1-ATR1-EDS1-PAD4, red) with retention time 4.5 min and synthesized standard pRib-ADP (black). Right: chromatograms of the compound from EDS1-PAD4 co-expressed with RPP1-ATR1 (RPP1-ATR1-EDS1-PAD4, green) with retention time 4.82 min and synthesized standard pRib-AMP (black). (**B**) Mirror image of MS/MS spectra from EDS1-PAD4 co-expressed with RPP1-ATR1 (top) and synthesized standard small molecules pRib-AMP (left bottom) and pRib-ADP (right bottom). (**C**) Omit electron density (2Fo-Fc) around pRib-ADP contoured at 1.5 sigma in the final structure of EDS1-PAD4. After modeling of pRib-AMP, a small portion of the density (color in blue) remains unoccupied. (**D**) Mass analysis of small molecules bound to EDS1-PAD4. Scheme showing the proposed routes of formation of the various ions of 558.0657(-) ion mass compound. Numbers in parentheses indicate the theoretical molecular weights of proposed ions.

**Fig. S3.**
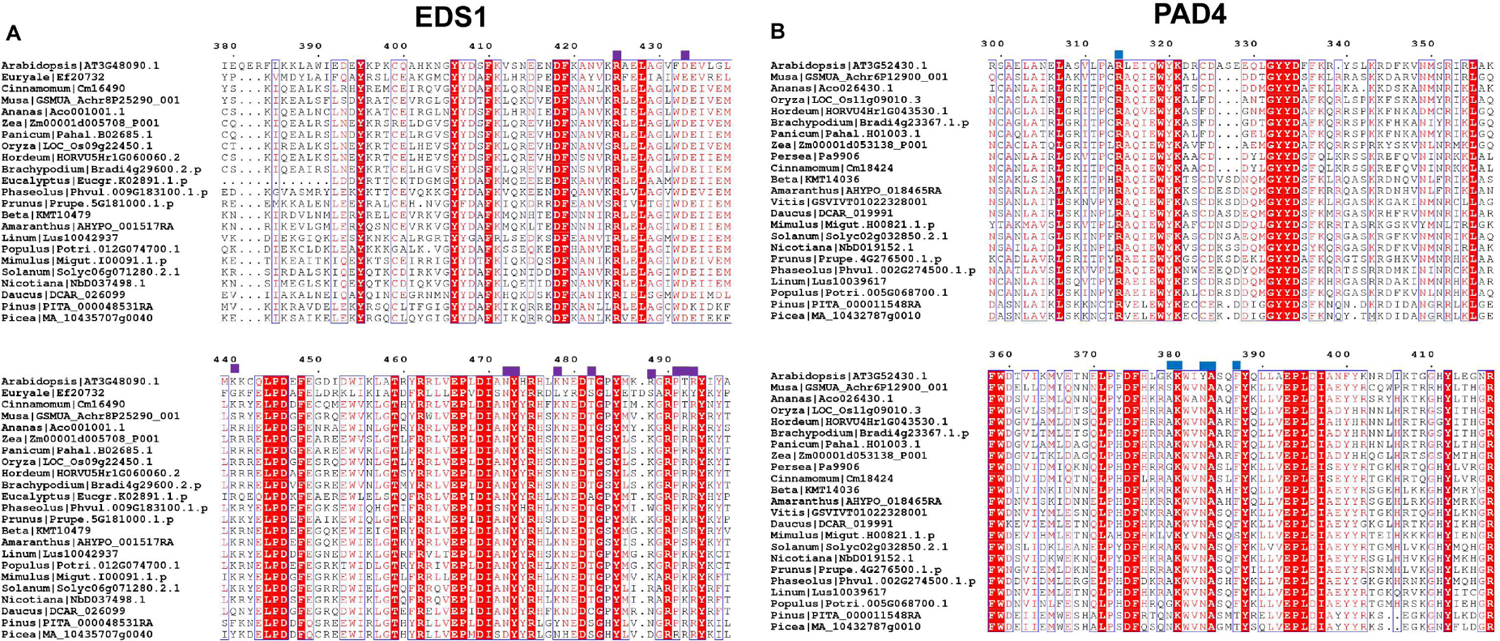
Sequence alignment of EDS1 and PAD4 from different plant species. (A) Sequence alignment of EDS1 from different plant species. Residues involved in recognition of pRib-ADP by *Arabidopsis* EDS1 marked with solid purple squares on the top. (B) Sequence alignment of PAD4 from different plant species. Residues involved in recognition of pRib-ADP by *Arabidopsis* PAD4 marked with solid blue squares on the top.

**Fig. S4.**
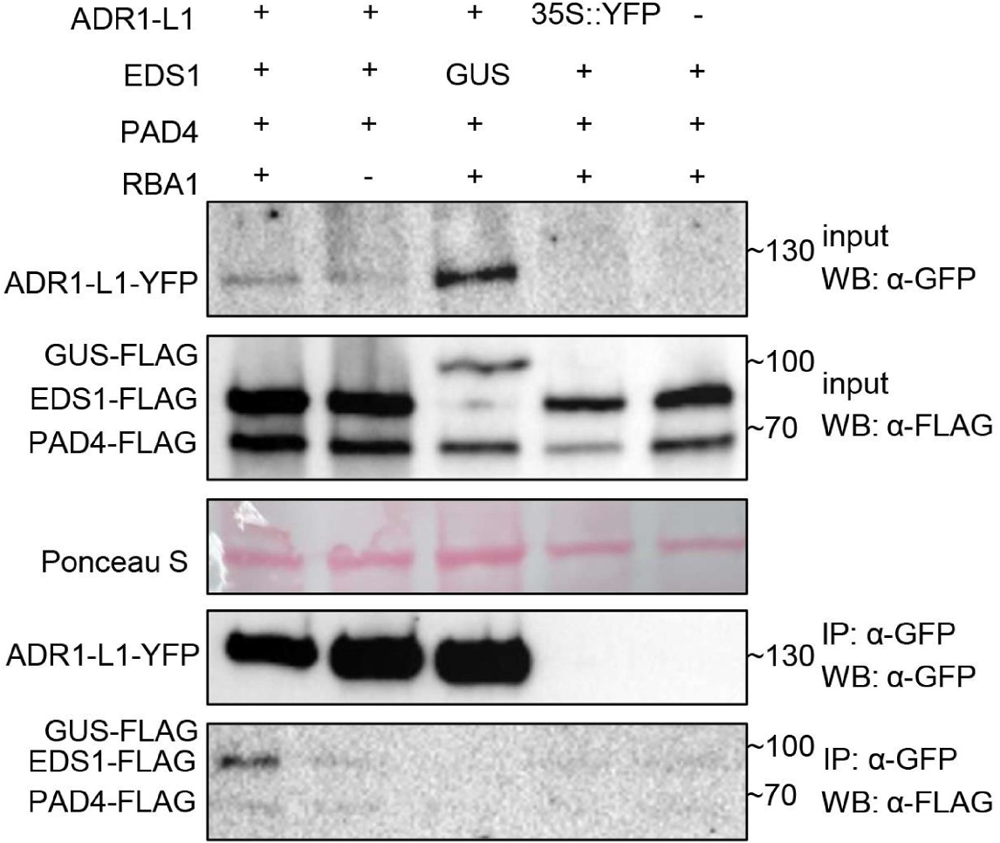
Expression of RBA1 promotes EDS1-PAD4 interaction with ADR1-L1 in *Nb* tobacco plants Co-immunoprecipitation (IP) assay followed by Western blotting to test for RBA1-triggered association of FLAG-tagged EDS1-PAD4 to ADR1-L1-YFP in *Nb* tobacco *epss* mutants. *Arabidopsis* EDS1-3xFLAG, PAD4-3xFLAG, GUS-3xFLAG, ADR1-L1-YFP and RBA1-3xmyc were expressed using Agrobacteria infiltration 48 h before sampling and protein extraction. Following IP of ADR1-L1-YFP using α-GFP agarose beads, input and IP fractions were probed with both α-GFP and α-FLAG antibodies. Expression of GUS-FLAG instead of EDS1-FLAG served as a negative control. The experiment was performed three times independently with similar results. WB: Western blotting, IP: immunoprecipitation.

**Fig. S5.**
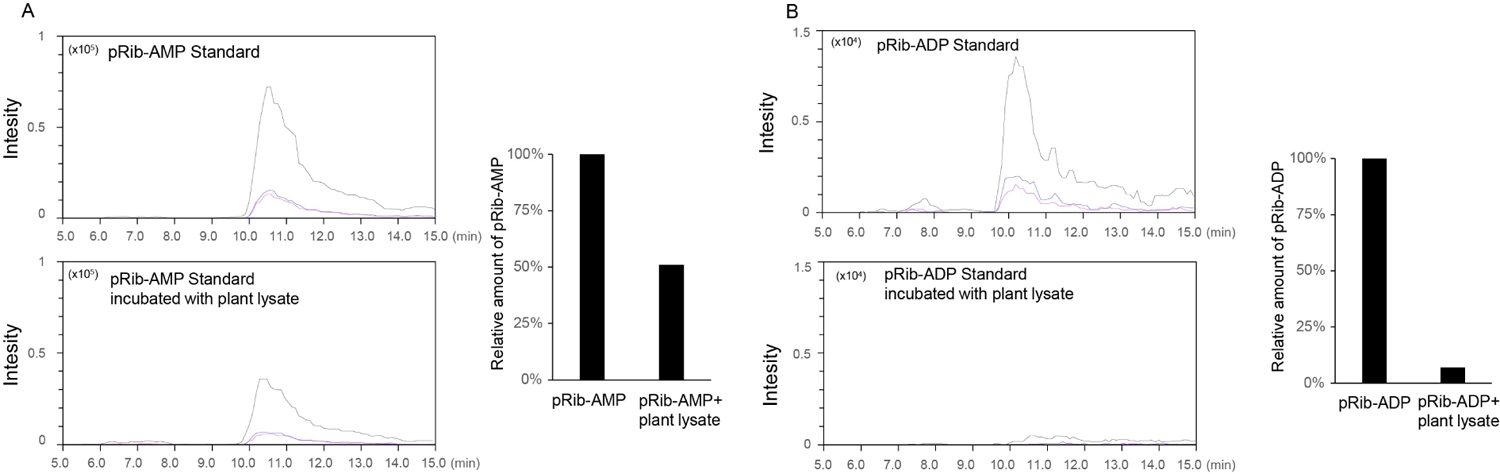
Degradation of pRib-AMP and pRib-ADP in lysate of *Nb* tobacco leaves. **(A)** The pRib-AMP standard (10 μM) was incubated with water and *Nb* tobacco leaf lysate for one hour at 4°C and the samples were analyzed by LC-MS. Black, red and blue lines indicate three different transitions 560.10>136.25, 560.10>348.00, 560.10>97.20 for pRib-AMP. The statistics shown in the right was based on the intensity of LC-MS results. The amount of pRib-AMP in water was normalized to 1.0. (**B**) The pRib-ADP standard (10 μM) was incubated with water and *Nb* tobacco leaf lysate for one h at 4°C and the samples were analyzed by LC-MS. Black, red and blue lines indicate three different transitions 640.10>136.25, 640.10>97.00, 640.10>428. 05 for pRib-ADP. The quantification shown on the right was based on the intensity of the peak in black. The amount of pRib-ADP in water was normalized to 1.0.

**Fig. S6.**
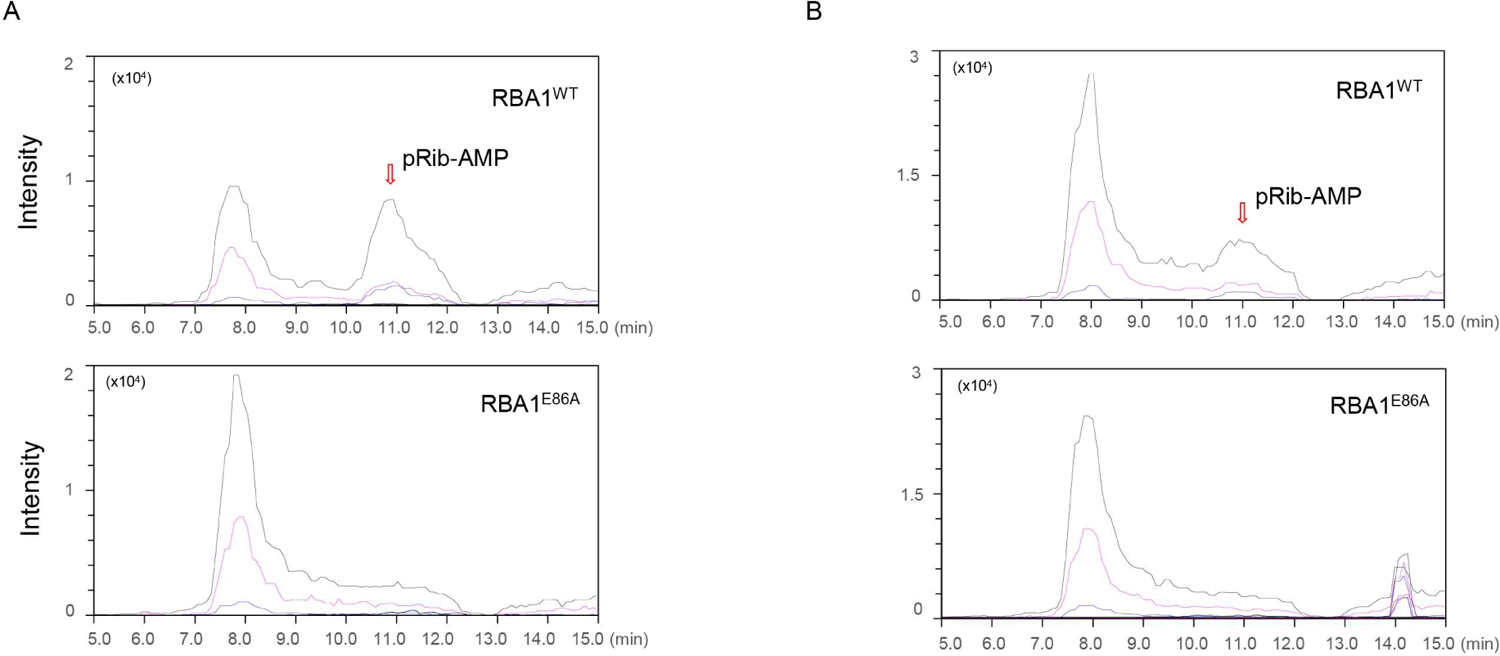
MRM analyses of pRib-AMP in *Nb* tobacco plants *Nb-epss* tobacco leaves expressing RBA1 or RBA1^E86A^ and insect cells expressing apo-EDS1-PAD4 complex were mixed and co-sonicated. EDS1-PAD4 complex protein in the lysate was purified through affinity chromatography and LC/MS was used to analyze the small molecules bound in the purified EDS1-PAD4 complex. Shown are representative chromatograms of pRib-AMP captured by EDS1-PAD4 in *Nb* tobacco expressing wild type *RBA1* (top) or the catalytic mutant *RBA1*^E86A^ (bottom). Black, purple and blue lines indicate three different transitions 560.10>136.20, 560.10>348.00, 560.10>97.20, respectively. “**A**” and “**B**” are the 2^nd^ and 3^rd^ replicates of the experiments.

**Fig. S7.**
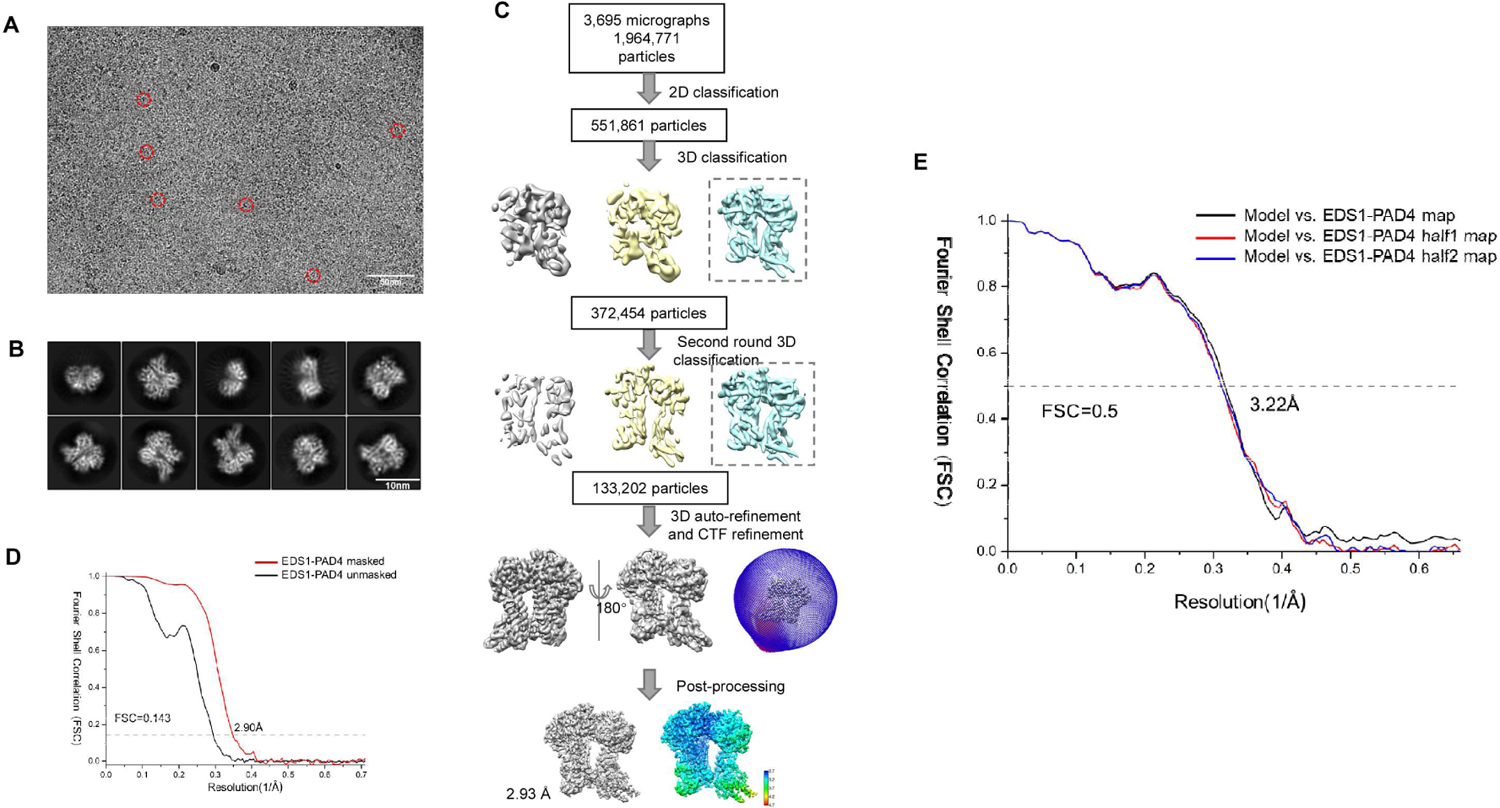
Cryo-EM reconstruction of the apo-EDS1-PAD4 complex. **(A)** A Representative cryo-EM micrograph of the apo-EDS1-PAD4. (**B**) Representative views of 2D class averages of the apo-EDS1-PAD4. (**C**) The cryo-EM image processing workflow. (**D**) FSC curves at 0.143 of the final reconstruction of the apo-EDS1-PAD4 complex unmasked (black) or masked (red). (**E**) FSC curves at 0.5 for model refined against the final map (black), the first half map (red) and the second half map (blue).

**Fig. S8.**
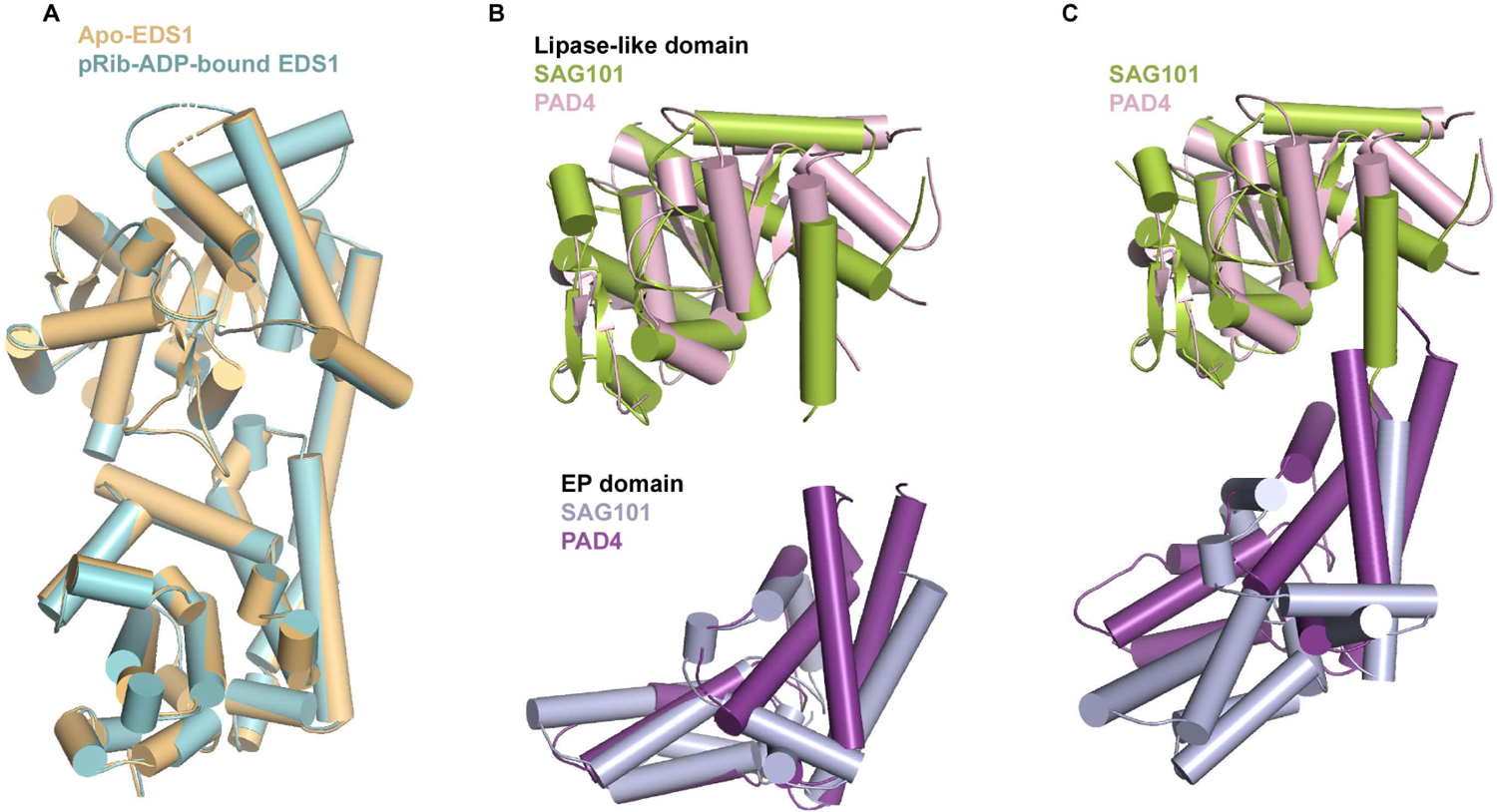
pRib-ADP binding to EDS1-PAD4 induces conformational changes in PAD4 EP domain. **(A)** Structural superposition of EDS1 in apo- and ligand-bound forms. EDS1 (light orange) from apo-EDS1-PAD4 was aligned with EDS1 (cyan) from pRib-ADP-bound form of EDS1-PAD4. (**B**) Structural comparison of the N-terminal lipase like domain (top) and C-terminal EP domain (bottom) of ligand-bound form of PAD4 with their counterparts of apo-form of SAG101. (**C**) Structural comparison between ligand bound form of PAD4 and apo-form of SAG101. The N-terminal lipase-like domain was used as the template for the structural alignment. Color codes for domains are the same those in (B).

**Fig. S9.**
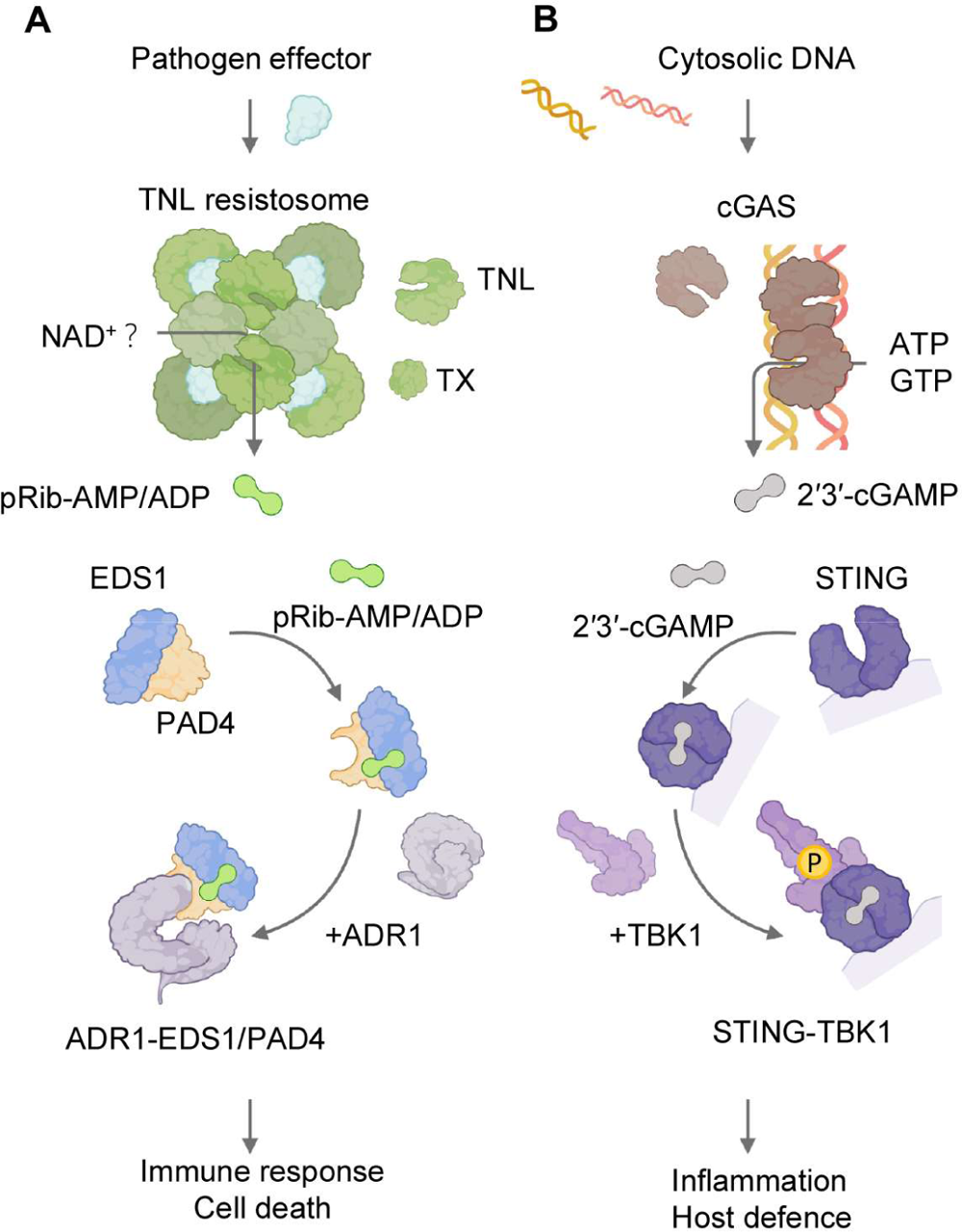
Working model on TIR-signaling. **(A)** pRib-AMP/ADP produced by pathogen effector-activated TNL or TIR-only (TX) proteins bind to EDS1-PAD4. Binding of either of the small molecules triggers a conformational change in the PAD4 EP domain to allosterically promote EDS1-PAD4 interaction with an ADR1 member. This may result in activation of the ADR1 resistosome and its Ca^2+^-channel activity, consequently initiating immune response. **(B)** 2’,3’-cGAMP-induced signaling in human.

**Table S1.** Crystallographic Data Collection and Refinement Statistics for the pRib-ADP-bound form EDS1-PAD4

| Data collection |  |
| --- | --- |
| Data set | EDS1-PAD4 |
| PDB ID | 7XEY |
| Wavelength (Å) | 0.979 |
| Resolution (Å) | 50-2.29 (2.33-2.29) |
| Space group | P 2<sub>1</sub> |
| Cell dimensions | 82.36 Å, 91.86 Å, 94.27 Å,<br>90.00, 111.79, 90.00 |
| Unique reflections | 57868 (2850) |
| Completeness (%) | 98.4 (98.0) |
| R<sub>meas</sub> (%) | 14.9 (78.3) |
| R<sub>pim</sub>(%) | 5.8 (35.3) |
| Redundancy | 6.4 (4.5) |
| Average *I*/σ(*I*) | 15.0 (2.0) |
| Statistics for Refinement |  |
| R<sub>work</sub> (%) | 18.8 (26.1) |
| R<sub>free</sub> (%) | 23.3 (32.5) |
| R.m.s.d. |  |
| Bond (degree) | 0.857 |
| Length (Å) | 0.006 |
| Ramachandran plot |  |
| Favored region (%) | 96.77 |
| Allowed region (%) | 2.69 |
| Outliers (%) | 0.54 |

**Table S2.**
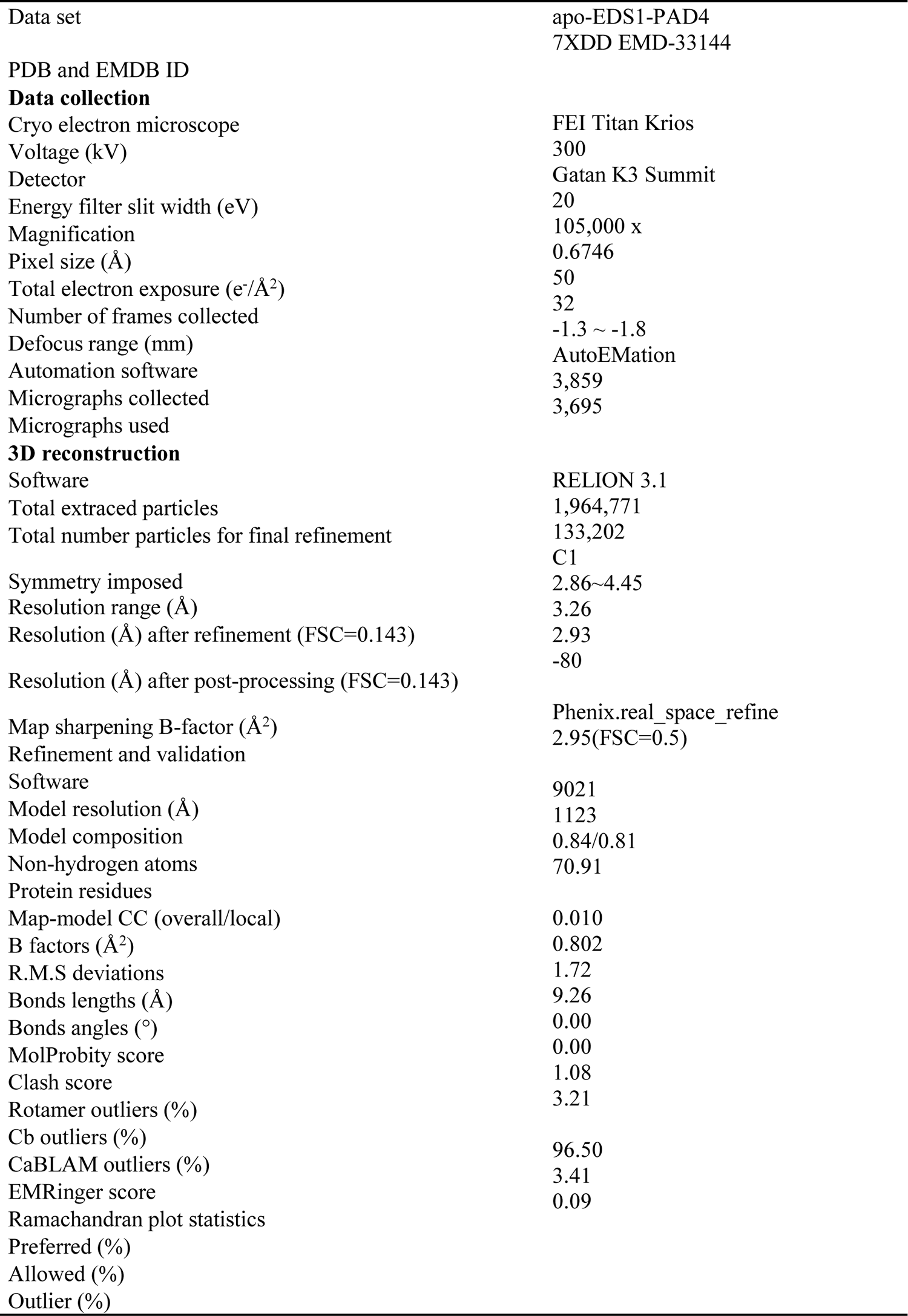
Cryo-EM statistics and model refinement for the apo-EDS1-PAD4

**Table S3.**
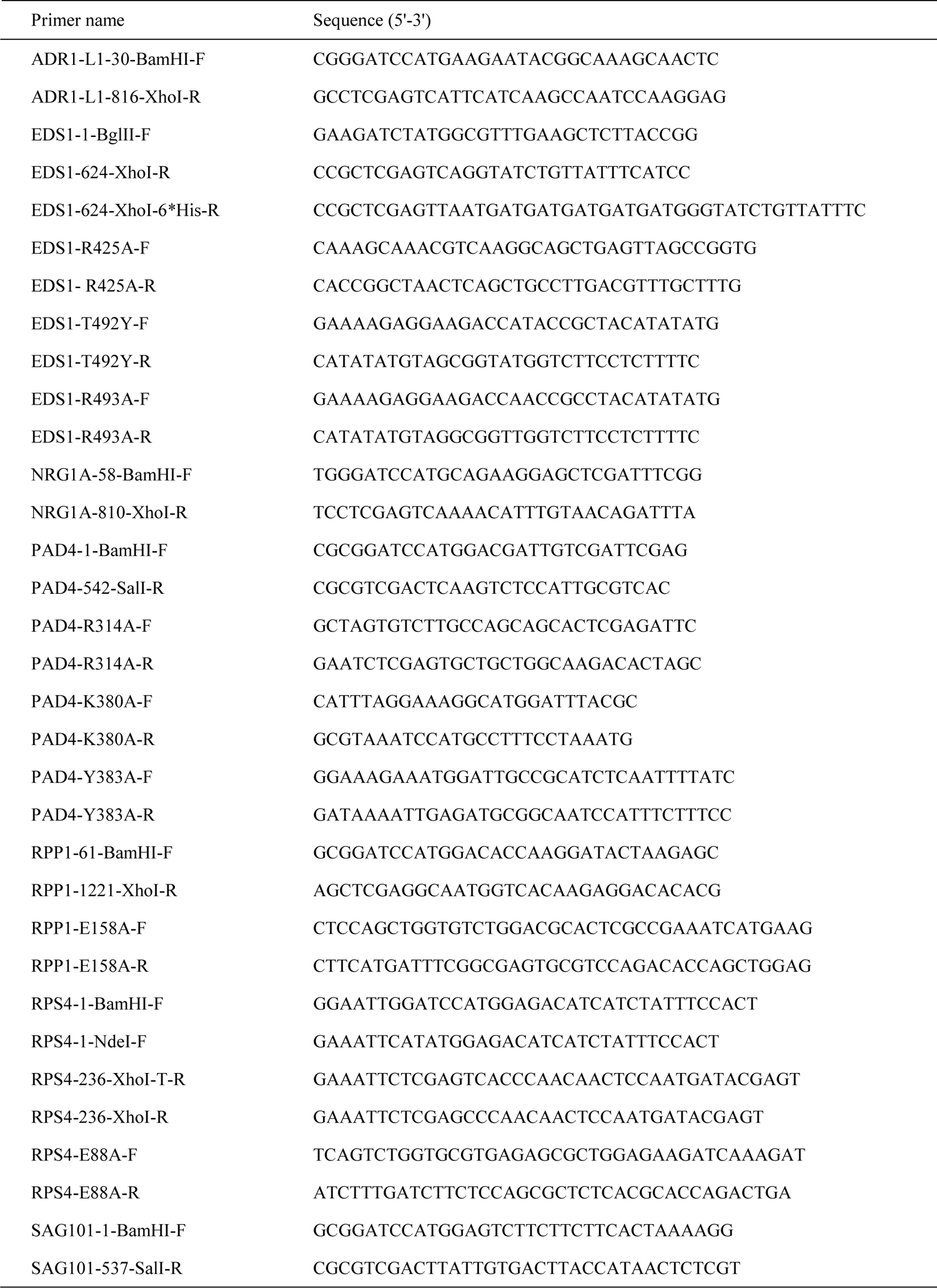
Oligonucleotides used in this study

### Data S1 Chemical synthesis of pRib-AMP/ADP

#### General Methods

All chemical reagents and solvents used in experiments were purchased from commercial suppliers and used without further purification. Silica gel (300−400 mesh) used for column chromatography and thin-layer chromatography (TLC) used to monitor progress of the reaction were purchased from Qingdao Haiyang Chemical Co., Ltd. ^1^H, ^31^P and ^13^C NMR spectrawere performed on the Bruker spectrometer (AV-400) at 400, 162 and 100 MHz, respectively. The mass spectra (MS, Thermo, LCQ, Fleet) were measured at the School of Pharmaceutical Sciences, Zhengzhou University

**Scheme 1.**
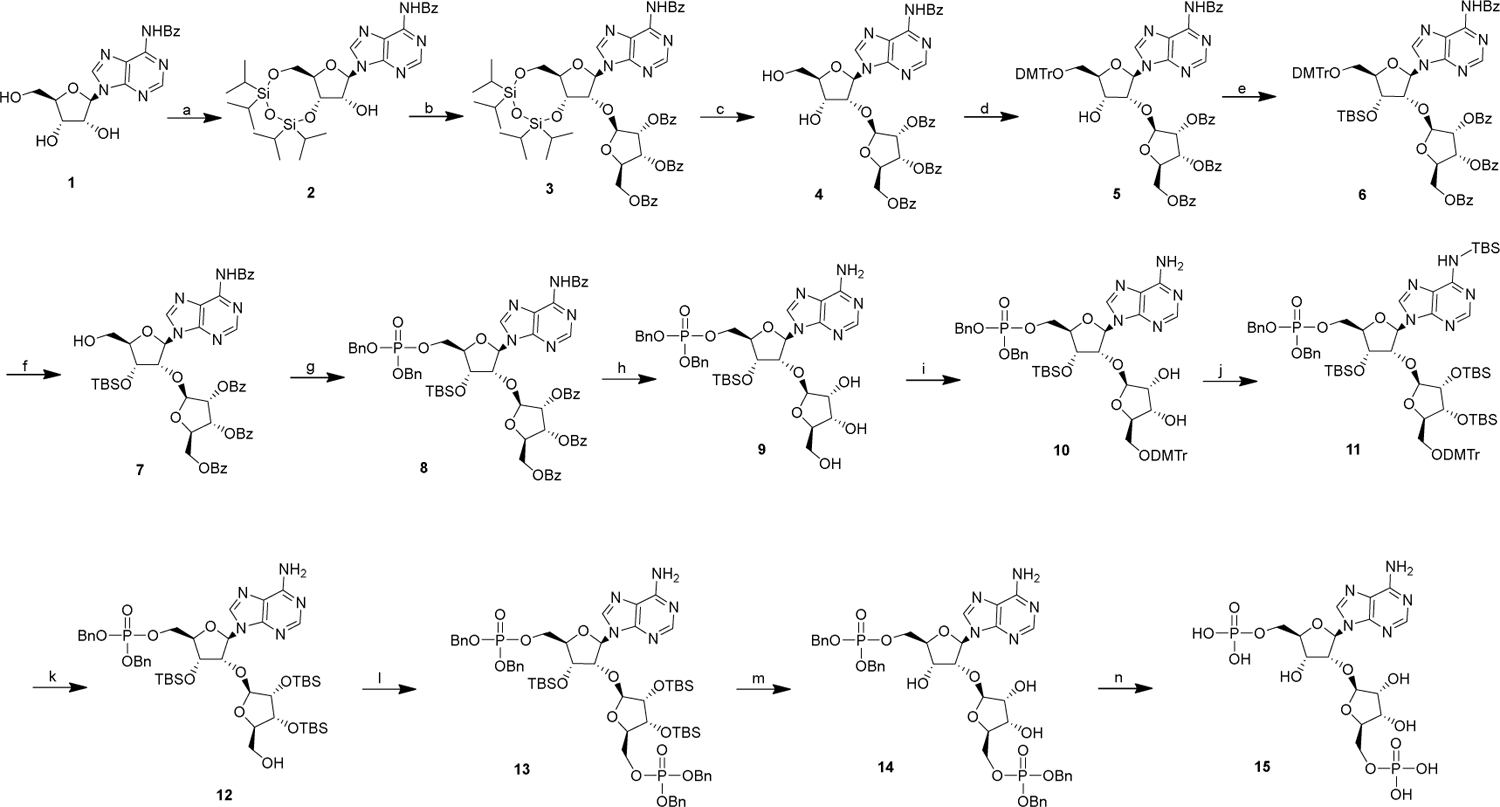
Synthesis of Compound **15**. Reagents and conditions: a) 1,3-dichloro-1,1,3,3-tetraisopropyldisiloxane, pyridine, overnight, 75%; b) beta-D-ribofuranose 1-acetate 2,3,5-tribenzoate, SnCl_4_, DCE, 30 h, 70%; c) TBAF, THF, 30 min, 80%; d) DMTrCl, pyridine, 2 h; e) TBSCl, imidazole, 2 h, two-step yield 61%; f) PTSA, CHCl_3_/MeOH, 10 min, 91%; g) dibenzyl diisopropylphosphoramidite, 4,5-dicyanoimidazole, TBHP, CH_2_Cl_2_/MeCN, 4.5 h, 71%; h) NH_3_/MeOH, 24 h, 80%; i) DMTrCl, pyridine, 2 h; j) TBSCl, imidazole, 2 h; k) *p*-toluenesulfonic acid monohydrate, CHCl_3_/MeOH, 10 min, three-step yield 58%; l) dibenzyl diisopropylphosphoramidite, 4,5-dicyanoimidazole, TBHP, CH_2_Cl_2_/MeCN, 4.5 h, 69%; m) TBAF, THF, 30 min; n) Pd/C, Et_3_N, H_2_, *t*-BuOH/H_2_O, two-step yield 50%.

**Compound 2:** To a stirring solution of **1** (30.00 g, 80.84 mmol) in pyridine (200 mL) at 0 °C was added 1,3-dichloro-1,1,3,3-tetraisopropyldisiloxane (26.77 g, 84.88 mmol), and the solution was stirred at room temperature for overnight. The crude reaction mixture was concentrated under reduced pressure and purified by silica gel column chromatography (CH_2_Cl_2_: MeOH = 95:5) to give compound **2** (37.00 g, 75%) as a white solid; ^1^H NMR (400MHz, CDCl_3_) *δ* 9.29 (s, 1H), 8.73 (s, 1H), 8.15 (s, 1H), 8.02 (d, *J* = 7.3 Hz, 2H), 7.58 (t, *J* = 7.2 Hz, 1H), 7.49 (t, *J* = 7.8 Hz, 2H), 6.02 (s, 1H), 5.08 (dd, *J* = 7.4, 5.6 Hz, 1H), 4.61 (d, *J* = 5.4 Hz, 1H), 4.15–4.01 (m, 3H), 3.45 (s, 1H), 1.12–0.99 (m, 28H) ppm; ^13^C NMR (100 MHz, CDCl_3_) *δ* 164.8, 152.8, 151.1, 149.8, 142.1, 133.8, 132.9, 128.9, 128.0, 123.7, 89.9, 82.3, 75.2, 70.7, 61.6, 17.54, 17.46, 17.44, 17.38, 17.2, 17.09, 17.07, 17.0, 13.4, 13.1, 12.9, 12.7 ppm; ESI-MS [M + H]^+^ for C_29_H_44_N_5_O_6_Si_2_, calcd 614.3, found 614.0.

**Compound 3:** To a stirring solution of beta-D-ribofuranose 1-acetate 2,3,5-tribenzoate (13.17 g, 26.10 mmol) in 1,2-dichloroethane (40 mL) at 0 °C was added tin tetrachloride (9.35 g, 35.88 mmol) under N_2_ atmosphere. After 10 min, a solution of compound **2** (1.62 g, 7.10 mmol) in 1,2-dichloroethane (30 mL) was added to the mixture and the reaction mixture was stirred at 0 °C for 30 h. The reaction mixture was quenched by the addition of the saturated aqueous NaHCO_3_, filtered through a pad of celite and washed with EtOAc. The organic layer was concentrated and purified via column chromatography (petroleum ether: EtOAc = 1:1) to afford compound **3** (12.10 g, 70%) as a white solid. ^1^H NMR (400MHz, CDCl_3_) *δ* 9.11 (s, 1H), 8.72 (s, 1H), 8.13 (s, 1H), 8.02–7.95 (m, 6H), 7.88 (d, *J* = 7.2 Hz, 2H), 7.60–7.56 (m, 2H), 7.53–7.47 (m, 4H), 7.42 (t, *J* = 8.0 Hz, 2H), 7.36–7.29 (m, 4H), 6.08 (s, 1H), 5.97 (dd, *J* = 6.5, 5.0 Hz, 1H), 5.89 (d, *J* = 4.9 Hz, 1H), 5.82 (s, 1H), 4.96 (dd, *J* = 9.2, 4.5 Hz, 1H), 4.88 (d, *J* = 4.6 Hz, 1H), 4.82–4.73 (m, 2H), 4.66 (dd, *J* = 11.2, 5.9 Hz, 1H), 4.18 (d, *J* = 13.3 Hz, 1H), 4.12 (d, *J* = 9.2 Hz, 1H), 4.01 (dd, *J* = 13.3, 2.4 Hz, 1H), 1.08–0.96 (m, 28H) ppm; ^13^C NMR (100 MHz, CDCl_3_) *δ* 166.2, 165.5, 165.1, 152.9, 150.9, 149.5, 142.0, 133.9, 133.62, 133.56, 133.3, 132.8, 129.9, 129.8, 129.7, 129.2, 129.0, 128.6, 128.50, 128.49, 128.0, 123.6, 105.8, 89.0, 81.5, 79.7, 78.7, 75.7, 72.7, 70.0, 65.5, 59.8, 17.5, 17.44, 17.39, 17.3, 17.2, 17.0, 16.9, 13.4, 13.0, 12.9, 12.7 ppm; ESI-MS [M + H]^+^ for C_55_H_64_N_5_O_13_Si_2_, calcd 1058.4, found 1058.2.

**Compound 4:** To a stirring solution of **3** (20.00 g, 18.91 mmol) in THF (100 mL) was added 1 M TBAF in THF (45 mL) at room temperature, and the mixture was stirred at the same temperature for 30 min. Water (40 mL) was added and the mixture was extracted with EtOAc (3 × 100 mL). The combined organic solution was dried over anhydrous Na_2_SO_4_, filtered, and concentrated. The residue was purified by silica gel column chromatography (CH_2_Cl_2_: MeOH = 95:5) to afford compound **4** (12.40 g, 80%) as a white solid. ^1^H NMR (400MHz, CDCl_3_) *δ* 9.57 (s, 1H), 8.79 (s, 1H), 8.29 (s, 1H), 8.03 (d, *J* = 7.4 Hz, 2H), 7.97 (d, *J* = 7.0 Hz, 2H), 7.86 (t, *J* = 7.0 Hz, 4H), 7.58 (t, *J* = 7.4 Hz, 1H), 7.53–7.47 (m, 5H), 7.36–7.30 (m, 6H), 6.15 (d, *J* = 7.2 Hz, 1H), 6.07 (d, *J* = 11.0 Hz, 1H), 5.74–5.68 (m, 2H), 5.25–5.22 (m, 2H), 4.65 (d, *J* = 4.4 Hz, 1H), 4.51–4.43 (m, 2H), 4.23 (s, 1H), 4.03 (dd, *J* = 11.7, 4.6 Hz, 1H), 3.94–3.85 (m, 2H), 3.71 (t, *J* = 1.6 Hz, 1H) ppm; ^13^C NMR (100 MHz, CDCl_3_) *δ* 166.0, 165.6, 165.5, 164.8, 152.3, 150.4, 144.1, 133.82, 133.76, 133.7, 133.5, 132.8, 129.83, 129.78, 129.7, 129.2, 128.8, 128.63, 128.60, 128.55, 128.52, 128.49, 128.08, 124.10, 106.5, 89.2, 87.3, 80.9, 80.0, 75.9, 72.4, 71. 5, 64.3, 63.1, 60.5 ppm; ESI-MS [M + H]^+^ for C_43_H_38_N_5_O_12_, calcd 816.3, found 816.0.

**Compound 6:** To a solution of compound **4** (10.00 g, 12.27 mmol) in pyridine (100 mL) was added DMTrCl (8.30 g, 24.53 mmol). The reaction mixture was stirred at room temperature for 2 h, after which MeOH (30 mL) was added. The mixture was rotoevaporated to dryness under reduced pressure to provide **5**. Unpurified **5** was dissolved in CH_2_Cl_2_ (100 mL), imidazole (1.30 g, 18.41 mmol) and TBSCl (2.70 g, 18.41 mmol) were added to the solution. The reaction mixture was stirred at room temperature for 2 h. Water (100 mL) was added, and the aqueous layer was extracted with CH_2_Cl_2_ (3 × 100 mL). The organic layer was dried over anhydrous Na_2_SO_4_ and concentrated. The residue was purified by silica gel column chromatography (petroleum ether: EtOAc = 2:1) to afford compound **6** (9.20 g, 61%) as a white solid. ^1^H NMR (400MHz, CDCl_3_) *δ* 9.14 (s, 1H), 8.69 (s, 1H), 8.14 (s, 1H), 8.03–7.95 (m, 6H), 7.84 (d, *J* = 7.0 Hz, 2H), 7.61–7.39 (m, 10H), 7.35–7.24 (m, 10H), 7.21–7.17 (m, 1H), 6.79 (dd, *J* = 9.0, 2.4 Hz, 4H), 6.21 (d, *J* = 4.8 Hz, 1H), 5.80–5.73 (m, 2H), 5.41 (s, 1H), 5.25 (t, *J* = 4.8 Hz, 1H), 4.63–4.55 (m, 2H), 4.39 (dd, *J* = 12.0, 4.1 Hz, 1H), 4.28 (dd, *J* = 12.0, 4.1 Hz, 1H), 4.22 (q, *J* = 4.6 Hz, 1H), 3.75 (s, 6H), 3.54 (dd, *J* = 11.0, 4.4 Hz, 1H), 3.29 (d, *J* = 11.0, 4.4 Hz, 1H), 0.85 (s, 9H), 0.11 (s, 3H), 0.02 (s, 3H) ppm; ^13^C NMR (100 MHz, CDCl_3_) *δ* 166.1, 165.3, 165.0, 164.6, 162.6, 158.6, 152.7, 151.4, 149.6, 144.5, 143.0, 135.7, 133.9, 133.7, 133.5, 133.3, 132.7, 130.1, 129.84, 129.77, 129.7, 129.4, 129.0, 128.9, 128.8, 128.6, 128.5, 128.4, 128.3, 127.92, 127.89, 127.0, 123.6, 113.2, 106.2, 87.9, 86.6, 84.3, 79.4, 79.3, 75.5, 72.1, 71.3, 64.7, 62.7, 55.3, 25.7, 18.1, −4.5, −4.9 ppm; ESI-MS [M + H]^+^ for C_70_H_70_N_5_O_14_Si, calcd 1232.5, found 1232.3.

**Compound 7:** The solution of *p*-toluenesulfonic acid monohydrate (1.20 g, 6.09 mmol) in MeOH (2 mL) was added to a solution of **6** (5.00 g, 4.06 mmol) in CHCl_3_ (20 mL) and the reaction mixture was stirred at room temperature for 10 min, whereupon saturated aqueous NaHCO_3_ (40 mL) was added to quench the reaction. The mixture was extracted with CH_2_Cl_2_ (3 × 100 mL). The organic layer was dried over anhydrous Na_2_SO_4_, filtered and concentrated. The residue was purified by silica gel column chromatography (CH_2_Cl_2_: MeOH = 95:5) to afford compound **7** (3.43 g, 91%) as a white solid. ^1^H NMR (400MHz, CDCl_3_) *δ* 9.32 (s, 1H), 8.78 (s, 1H), 8.10 (s, 1H), 8.04–8.00 (m, 4H), 7.94–7.91 (m, 2H), 7.83–7.80 (m, 2H), 7.61–7.46 (m, 6H), 7.38 (t, *J* = 7.8 Hz, 4H), 7.29 (t, *J* = 7.8 Hz, 2H), 6.16 (d, *J* = 10.1 Hz, 1H), 6.04 (d, *J* = 7.6 Hz, 1H), 5.67 (t, *J* = 5.3 Hz, 1H), 5.60 (dd, *J* = 5.1, 2.4 Hz, 1H), 5.17–5.14 (m, 2H), 4.62 (d, *J* = 4.8 Hz, 1H), 4.35 (q, *J* = 4.9 Hz, 1H), 4.28 (dd, *J* = 11.8, 4.2 Hz, 1H), 4.21 (s, 1H), 4.06 (dd, *J* = 11.8, 4.2 Hz, 1H), 3.96 (d, *J* = 13.2 Hz, 1H), 3.74 (d, *J* = 11.6 Hz, 1H), 0.88 (s, 9H), 0.08 (s, 3H), 0.07 (s, 3H) ppm; ^13^C NMR (100 MHz, CDCl_3_) *δ* 166.1, 165.3, 165.0, 164.6, 152.2, 150.47, 150.46, 144.2, 133.8, 133.7, 133.6, 133.5, 132.9, 129.83, 129.80, 129.78, 129.3, 128.9, 128.82, 128.76, 128.7, 128.6, 128.5, 128.0, 124.4, 106.1, 89.7, 89.3, 80.6, 79.5, 75.1, 72.7, 72.1, 64.6, 63.1, 25.8, 18.3, −4.6, −4.8 ppm; ESI-MS [M + H]^+^ for C_49_H_52_N_2_O_12_Si, calcd 930.3, found 930.1.

**Compound 8:** A round-bottom flask was charged with compound **7** (2.00 g, 2.15 mmol) and dibenzyl diisopropylphosphoramidite (890 mg, 2.58 mmol), anhydrous MeCN (30 mL) was added and rotoevaporated to dryness under reduced pressure for three times. The mixture was dissolved in CH_2_Cl_2_ (20 mL), whereupon the suspension of 4,5-dicyanoimidazole (310 mg, 2.58 mmol) in anhydrous MeCN (3 mL) was added under N_2_ atmosphere. The reaction was stirred at room temperature for 1.5 h; then 5.5 M TBHP (2 mL) was added to the mixture, which was kept stirring over a period of 3 h and then saturated aqueous NaHCO_3_ (10 mL) was added. The mixture was extracted with CH_2_Cl_2_ (3 × 50 mL). The organic layer was dried over anhydrous Na_2_SO_4_, filtered and concentrated. The residue was purified by silica gel column chromatography (CH_2_Cl_2_: MeOH = 95:5) to afford compound **8** (1.82 g, 71%) as a white solid. ^1^H NMR (400MHz, CDCl_3_) *δ* 9.11 (s, 1H), 8.69 (s, 1H), 8.11 (s, 1H), 7.99–7.93 (m, 6H), 7.83– 7.80 (m, 2H), 7.58–7.52 (m, 2H), 7.50–7.42 (m, 4H), 7.38 (t, *J* = 7.9 Hz, 2H), 7.33–7.22 (m, 14H), 6.16 (d, *J* = 4.4 Hz, 1H), 5.76 (dd, *J* = 6.5, 4.89 Hz, 1H), 5.70 (dd, *J* = 4.9, 1.4 Hz, 1H), 5.37 (d, *J* = 1.4 Hz, 1H), 5.10 (t, *J* = 4.7 Hz, 1H), 4.98–4.90 (m, 4H), 4.61 (t, *J* = 4.7 Hz, 1H), 4.55–4.51 (m, 1H), 4.42 (dd, *J* = 12.0, 4.0 Hz, 1H), 4.29–4.24 (m, 2H), 4.13–4.05 (m, 2H), 0.85 (s, 9H), 0.10 (s, 3H), 0.07 (s, 3H) ppm; ^31^P NMR (162 MHz, CDCl_3_) *δ* −0.92; ESI-MS [M + H]^+^ for C_63_H_65_N_5_O_15_PSi, calcd 1190.4, found 1190.0.

**Compound 9:** A solution of **8** (5.00 g, 4.20 mmol) in NH_3_/MeOH (40 mL) was stirred for 24 h at room temperature. The mixture was concentrated under reduced pressure and purified by silica gel column chromatography (CH_2_Cl_2_: MeOH = 20:1) to afford compound **9** (2.60 g, 80%) as a white solid. ^1^H NMR (400MHz, CD_3_OD) *δ* 8.25 (s, 1H), 8.16 (s, 1H), 7.31–7.24 (m,10H), 6.28–6.26 (m, 1H), 4.99–4.92 (m, 5H), 4.78 (t, *J* = 4.6 Hz, 1H), 4.71 (t, *J* = 4.6 Hz, 1H), 4.35–4.30 (m, 1H), 4.20–4.10 (m, 3H), 4.02–3.99 (m, 1H), 3.89 (m, 1H), 3.67–3.60 (m, 1H), 3.47–3.41 (m, 1H), 0.93 (s, 9H), 0.15 (s, 3H), 0.13 (s, 3H); ^31^P NMR (162 MHz, CD_3_OD) *δ* −1.47; ESI-MS [M + H]^+^ for C_35_H_49_N_5_O_11_PSi, calcd 774.3, found 774.1.

**Compound 11:** To a solution of product **9** (2.00 g, 2.58 mmol) in pyridine (40 mL) was added DMTrCl (1.75 g, 5.17 mmol). The reaction mixture was stirred at room temperature for 2 h, after which MeOH (30 mL) was added. The mixture was rotoevaporated to dryness under reduced pressure to provide **10**. Unpurified **10** was dissolved in CH_2_Cl_2_ (40 mL), to which was added imidazole (1.30 g, 18.41 mmol) followed by TBSCl (2.70 g, 18.41 mmol), and the reaction mixture was stirred at room temperature for 2 h. Water (50 mL) was added, and the aqueous layer was extracted with CH_2_Cl_2_ (3 × 100 mL). The organic layer was dried over anhydrous Na_2_SO_4_ and concentrated to afford compound **11**. To a stirring solution of **11** in CHCl_3_ (30 mL) was added the solution of *p*-toluenesulfonic acid monohydrate (1.20 g, 6.09 mmol) in MeOH (5 mL), and the reaction mixture was stirred at room temperature for 10 min, whereupon saturated aqueous NaHCO_3_ (40 mL) was added to quench the reaction. The mixture was extracted with CH_2_Cl_2_ (3 × 50 mL). The organic layer was dried over anhydrous Na_2_SO_4_, filtered and concentrated. The residue was purified by silica gel column chromatography (CH_2_Cl_2_: MeOH = 95:5) to give compound **12**(1.93 g, 58%) as a white solid. ^1^H NMR (400MHz, CDCl_3_) *δ* 8.28 (s, 1H), 8.22 (s, 1H), 7.29–7.17 (m, 10H), 6.30 (d, *J* = 1.3 Hz, 1H), 6.08 (s, 1H), 5.10–4.97 (m, 4H), 4.87 (s, 1H), 4.42–4.31 (m, 3H), 4.22 (d, *J* = 4.4 Hz, 1H), 4.12–4.05 (m, 2H), 3.98 (d, *J* = 3.8 Hz, 1H), 3.94–3.89 (m,2 H), 3.59 (d, *J* = 12.4 Hz, 1H), 0.87 (s, 9H), 0.82 (s, 9H), 0.81 (s, 9H), 0.13 (s, 3H), 0.07 (s, 3H), 0.04 (s, 3H), 0.02 (s, 3H), 0.00 (s, 3H), −0.01 (s, 3H) ppm, ^31^P NMR (162 MHz, CDCl_3_) *δ* −1.76; ESI-MS [M + H]^+^ for C_47_H_77_N_5_O_11_PSi_3_, calcd 1002.5, found 1002.2.

**Compound 13:** The compound **13** was synthesized from **12** (35 mg, 0.11 mmol) by the procedure described for the synthesis of **8** (1.82 g, 69%). ^1^H NMR (400MHz, CDCl_3_) *δ* 8.14 (s, 1H), 8.10 (s, 1H), 7.26–7.12 (m, 20H), 6.22 (d, *J* = 2.6 Hz, 1H), 5.86 (s, 2H), 5.00–4.93 (m, 4H), 4.86–4.77 (m, 7H), 4.25–3.89 (m, 8H), 0.86 (s, 9H), 0.80 (s, 9H), 0.79 (s, 9H), 0.05 (s, 6H), 0.00–-0.02 (m, 12H) ppm, ^31^P NMR (162 MHz, CDCl_3_) *δ* −0.51, −1.23; ESI-MS [M + H]^+^ for C_61_H_90_N_5_O_14_P_2_Si_3_, calcd 1262.5, found 1262.2.

**Compound 15:** To a flask containing **13** (200 mg, 0.16 mmol) was added THF (10 mL) followed by 1 M TBAF in THF (0.55 mL) and the mixture was stirred for 30 min. Water (20 mL) was added and the mixture was extracted with CH_2_Cl_2_ (3 × 50 mL). The combined organic solution was dried over anhydrous Na_2_SO_4_, filtered. The filtrate was rotoevaporated to provide compound **14.** The crude product **14** prepared above was dissolved in *t*-BuOH (20 mL) to which was added H_2_O (2 mL) followed by Et_3_N (0.05 mL) and Pd/C (20 mg), then the reaction was stirred for overnight under H_2_ atmosphere. The mixture was filtered through a pad of celite and the filtrate was evaporated under high vacuum to afford a triethylamine salt of compound **15** (45 mg, 50%); HRMS(ESI)calcd. for C_15_H_23_N_5_O_14_P_2_ ([M + H]^+^):560.0790, found: 560.0787.

**Scheme 2.**
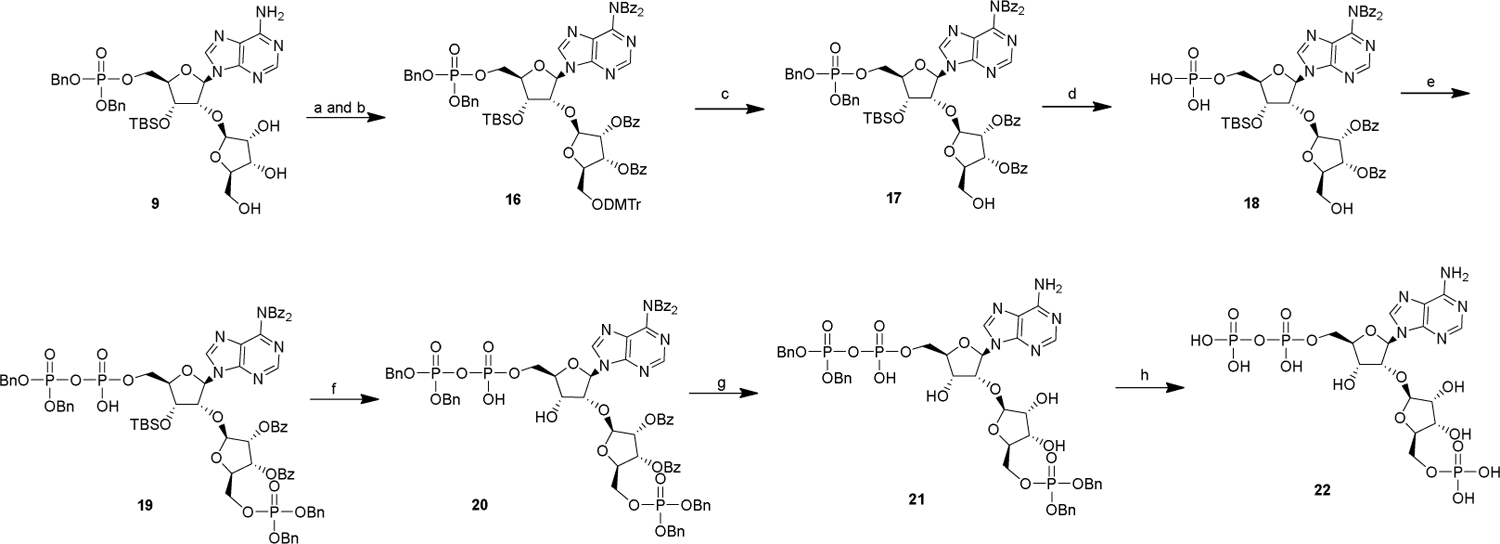
Synthesis of Compound **22**. Reagents and conditions: a) DMTrCl, pyridine, 2 h; b) DMAP, benzoyl chloride, Et_3_N, CH_2_Cl_2_, pyridine, 5 h, two-step yield 62%; c) *p*-toluenesulfonic acid monohydrate, CHCl_3_/MeOH, 10 min, 90%; d) Pd/C, H_2_, *t*-BuOH; e) dibenzyl diisopropylphosphoramidite, 4,5-dicyanoimidazole, TBHP, CH_2_Cl_2_/MeCN, 4.5 h, two-step yield 43%; f) TBAF, THF, 30 min; g) NH_3_/MeOH, 24 h; h) Pd/C, Et_3_N, H_2_, *t*-BuOH /H_2_O, three-step yield 25%.

**Compound 16:** To a solution of product **9** (2.00 g, 2.58 mmol) in pyridine (40 mL) was added DMTrCl (1.75 g, 5.17 mmol). The reaction mixture was stirred at room temperature for 2 h, after which MeOH (30 mL) was added. The mixture was rotoevaporated to dryness under reduced pressure to provide **10**. Unpurified **10** was dissolved in CH_2_Cl_2_ (20 mL) at room temperature, to which was added Et_3_N (2.15 mL,15.48 mmol)、DMAP (190 mg, 1.55 mmol)、benzoyl chloride (2.18 g, 15.48 mmol), and then the mixture was stirred for 5 h at room temperature. Water (20 mL) was added, and the aqueous phase was extracted with CH_2_Cl_2_ (3 × 30 mL). The organic layer was dried over anhydrous Na_2_SO_4_, filtered and concentrated. The residue was purified by silica gel column chromatography (petroleum ether: EtOAc = 2:1) to give compound **16** (2.39 g, 62%) as a white solid. ^1^H NMR (400MHz, CDCl_3_) *δ* 8.34 (s, 1H), 7.95–7.74 (m, 10H), 7.47–7.42 (m, 2H), 7.37–7.16 (m, 25H), 7.12–7.02 (m, 3H), 6.66–6.63 (m, 4H), 6.06 (d, *J* = 2.6 Hz, 1H), 5.79 (t, *J* = 5.2 Hz, 1H), 5.72 (dd, *J* = 4.9, 2.2 Hz, 1H), 5.29 (d, *J* = 2.1 Hz, 1H), 4.89 (s, 2H), 4.87 (s, 2H), 4.71 (dd, *J* = 5.3, 2.6 Hz, 1H), 4.62 (t, *J* = 6.0 Hz, 1H), 4.32 (q, *J* = 4.3 Hz, 1H), 4.31–4.24 (m, 1H), 4.10–4.05 (m, 2H), 3.59 (s, 3H), 3.58 (s, 3H), 3.37 (dd, *J* = 10.4, 4.2 Hz, 1H), 3.31 (dd, *J* = 10.4, 4.2 Hz, 1H), 0.76 (s, 9H), 0.00 (s, 3H), −0.01 (s, 3H) ppm; ^31^P NMR (162 MHz, CDCl_3_) *δ* −0.86; ESI-MS [M + Na]^+^ for C_84_H_82_N_5_NaO_17_PSi, calcd 1514.5, found 1514.2.

**Compound 17:** To a stirring solution of **16** (600 mg, 0.40 mmol) in CHCl_3_ (30 mL) was added the solution of *p*-toluenesulfonic acid monohydrate (100 mg, 0.52 mmol) in MeOH (5 mL), and the solution was stirred at room temperature for 10 min, whereupon saturated aqueous NaHCO_3_ (40 mL) was added to quench the reaction. The mixture was extracted with CH_2_Cl_2_ (3 × 50 mL). The organic layer was dried over anhydrous Na_2_SO_4_, filtered. The filtrate was evaporated in vacuo and subjected to column chromatography (CH_2_Cl_2_: MeOH = 95:5) yielding compound **17** (430 mg, 90%) as a white solid. ^1^H NMR (400MHz, CDCl_3_) *δ* 8.54 (s, 1H), 8.42 (s, 1H), 7.96–7.77 (m, 9H), 7.50–7.17 (m, 25H), 6.23 (d, *J* = 2.9 Hz, 1H), 5.75 (dd, *J* = 7.6, 4.8 Hz, 1H), 5.68 (d, *J* = 4.8 Hz, 1H), 5.01–4.91 (m, 4H), 4.57(dd, *J* = 4.6, 3.8 Hz, 1H), 4.37–4.27 (m, 1H), 4.18–4.14 (m, 1H), 4.10–4.06 (m, 1H), 3.73 (dd, *J* = 12.8, 2.4 Hz, 1H), 3.50 (dd, *J* = 12.8, 3.8 Hz, 1H), 0.75 (s, 9H), 0.00 (s, 3H), −0.02 (s, 3H) ppm, ^31^P NMR (162 MHz, CDCl_3_) *δ* −0.52; ESI-MS [M + H]^+^ for C_63_H_65_N_5_O_15_PSi, calcd 1190.4, found 1190.4.

**Compound 19:** To a solution of **17** (400 mg, 0.34 mmol) in *t*-BuOH (20 mL) was added Pd/C (20 mg) and the reaction was stirred for overnight under H_2_ atmosphere. The suspension was filtered through a pad of celite and the filtrate was evaporated under low pressure to afford compound **18**. A round-bottom flask was charged with compound **18** and dibenzyl diisopropylphosphoramidite (290 mg, 0.84 mmol), anhydrous MeCN (30 mL) was added and rotoevaporated to dryness under reduced pressure for three times. The mixture was dissolved in CH_2_Cl_2_ (20 mL), whereupon the suspension of 4,5-dicyanoimidazole (100 mg, 0.84 mmol) in anhydrous MeCN (3 mL) was added under N_2_ atmosphere. The reaction mixture was stirred at room temperature for 1.5 h; the 5.5 M TBHP (0.6 mL) was added to the mixture, which was kept stirring over a period of 3 h and then saturated aqueous NaHCO_3_ (15 mL) was added. The mixture was extracted with CH_2_Cl_2_ (3 × 30 mL). The organic layer was dried over anhydrous Na_2_SO_4_, filtered and concentrated. The residue was purified by silica gel column chromatography (CH_2_Cl_2_: MeOH = 95:5) to afford compound **19** (225 mg, 43%) as a white solid. ^1^H NMR (400MHz, CDCl_3_) *δ* 8.60 (s, 1H), 7.91 (d, *J* = 7.8 Hz, 2H), 7.82 (d, *J* = 7.8 Hz, 2H), 7.65 (d, *J* = 7.6 Hz, 4H), 7.56 (t, *J* = 7.4 Hz, 1H), 7.49 (t, *J* = 7.4 Hz, 1H), 7.38 (t, *J* = 7.7 Hz, 2H), 7.31–7.08 (m, 29H), 6.06–5.83 (m, 1H), 5.68–5.45 (m, 3H), 5.16–4.90 (m, 8H), 4.74–4.47 (m, 2H), 4.32–3.66 (m, 6H), 0.74 (s, 9H), −0.05 (s, 6H); ^31^P NMR (162 MHz, CDCl_3_) δ − 1.68, −10.90, −12.37; ESI-MS [M - H]^-^ for C_77_H_77_N_5_O_21_P_3_Si, calcd1528.4, found 1528.4.

**Compound 22:** To a flask containing **19** (200 mg, 0.13 mmol) was added THF (10 mL) followed by 1 M TBAF in THF (0.15 mL) and the mixture was stirred for 30 min. Water (20 mL) was added and the mixture was extracted with CH_2_Cl_2_ (3 × 50 mL). The combined organic solution was dried over anhydrous Na_2_SO_4_, filtered. The filtrate was rotoevaporated to provide compound **20.** Unpurified **20** was dissolved in NH_3_/MeOH (20 mL), which was stirred for 24 h at room temperature. After completion of the reaction as indicated by mass spectrometry, the reaction was evaporated under reduced pressure to give **21** that was used directly in the next step without purification. The crude product **21** was dissolved in *t*-BuOH (20 mL) to which was added H_2_O (2 mL) followed by Et_3_N (0.05 mL) and Pd/C (20 mg), then the reaction was stirred for overnight under H_2_ atmosphere. After completion of the reaction as indicated by mass spectrometry, the mixture was filtered through a pad of celite and the filtrate was evaporated under high vacuum to afford a triethylamine salt of compound **22** (20 mg, 25%); HRMS( ESI) calcd. For C_15_H_24_N_5_O_17_P_3_([M + H]^+^): 640.0453, found (640.0454.

#### NMR spectra of the compounds

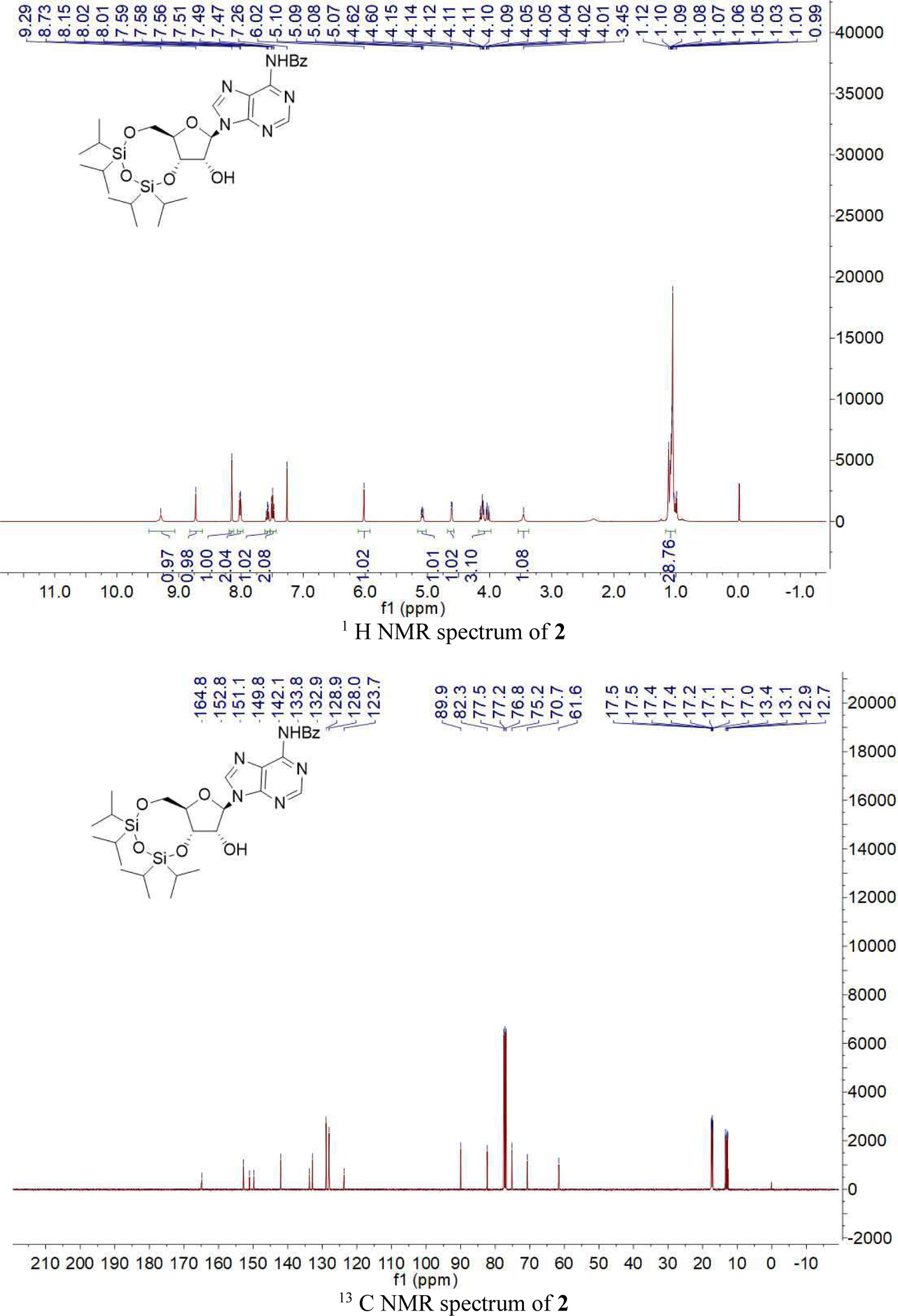
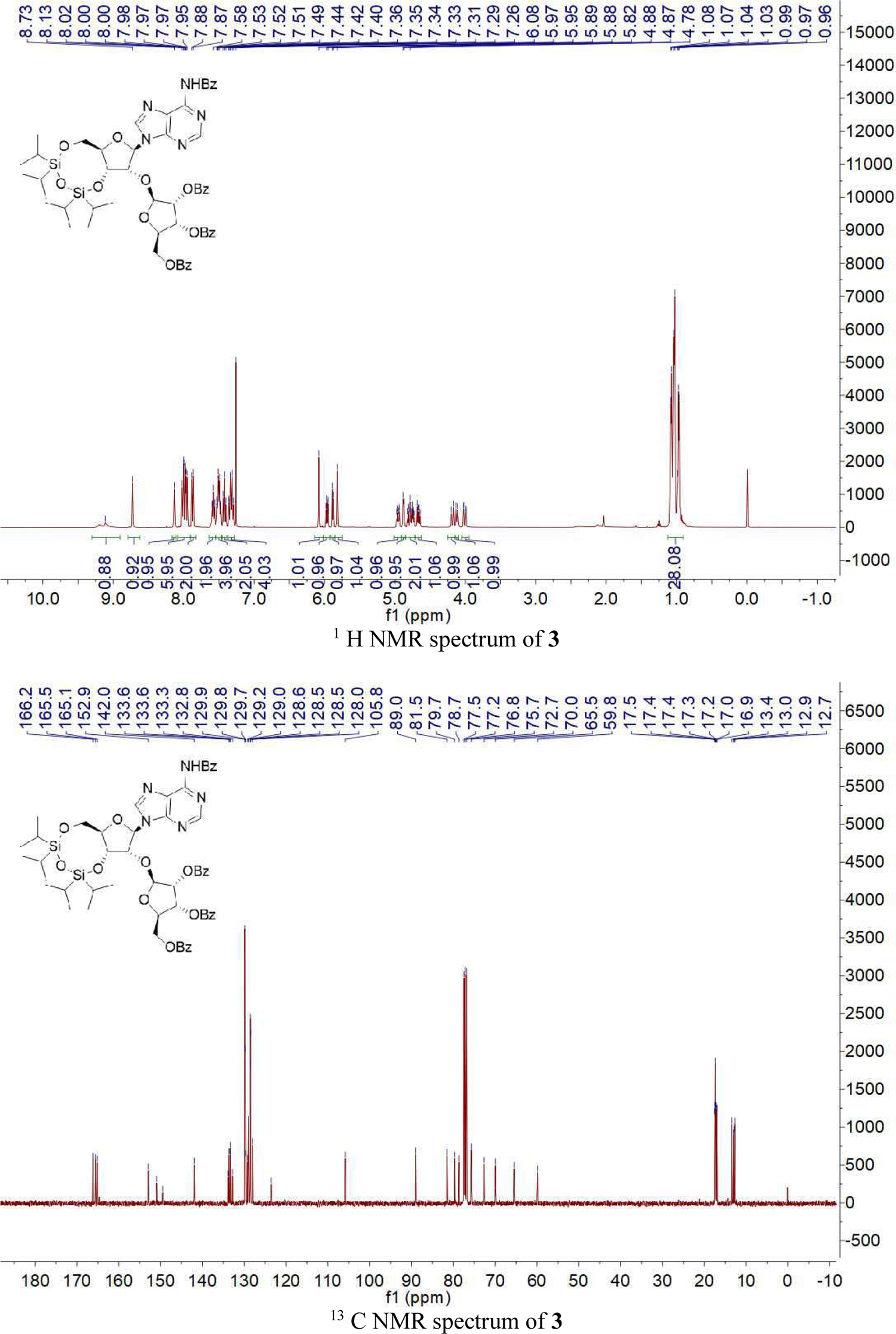
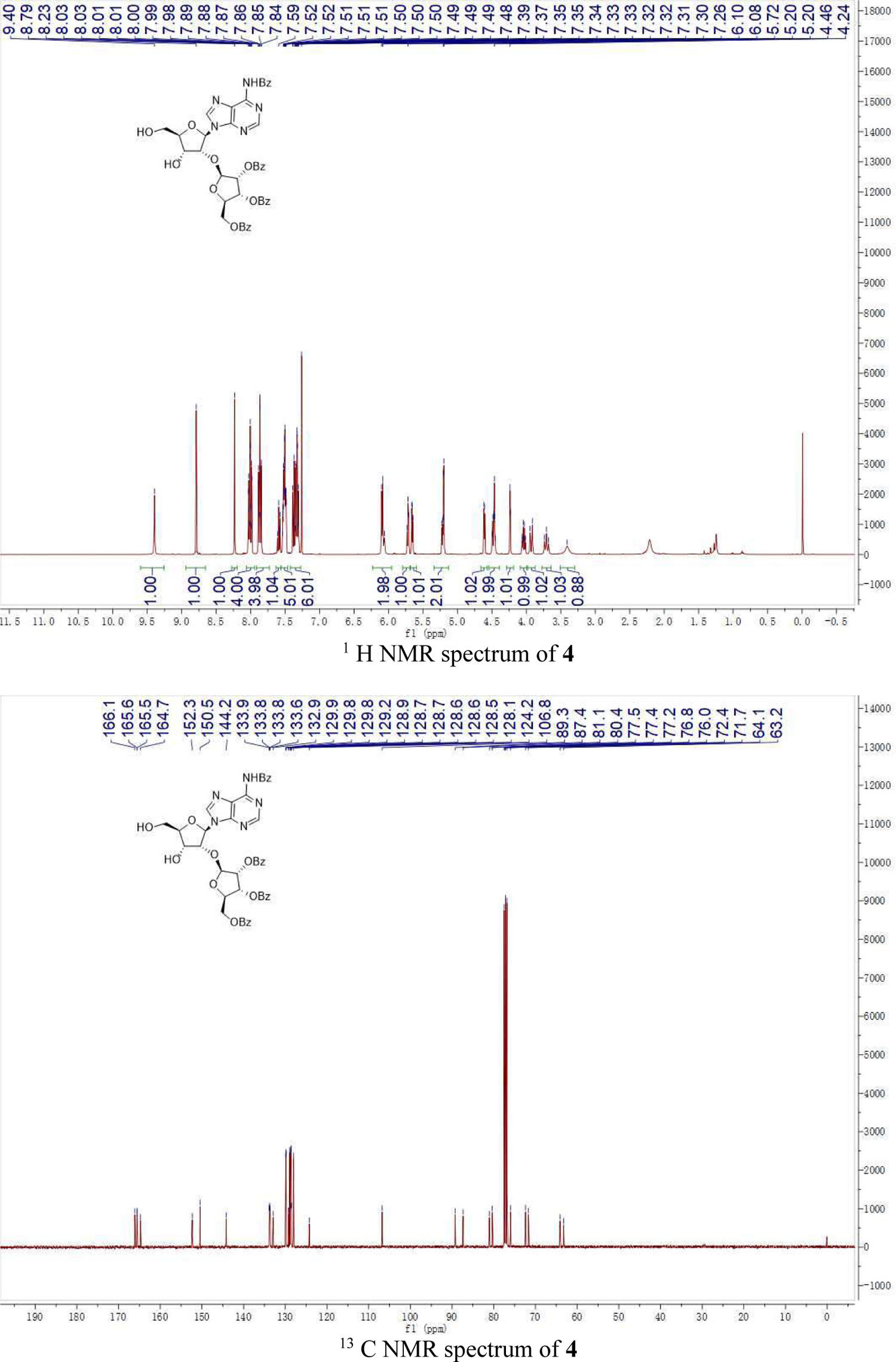
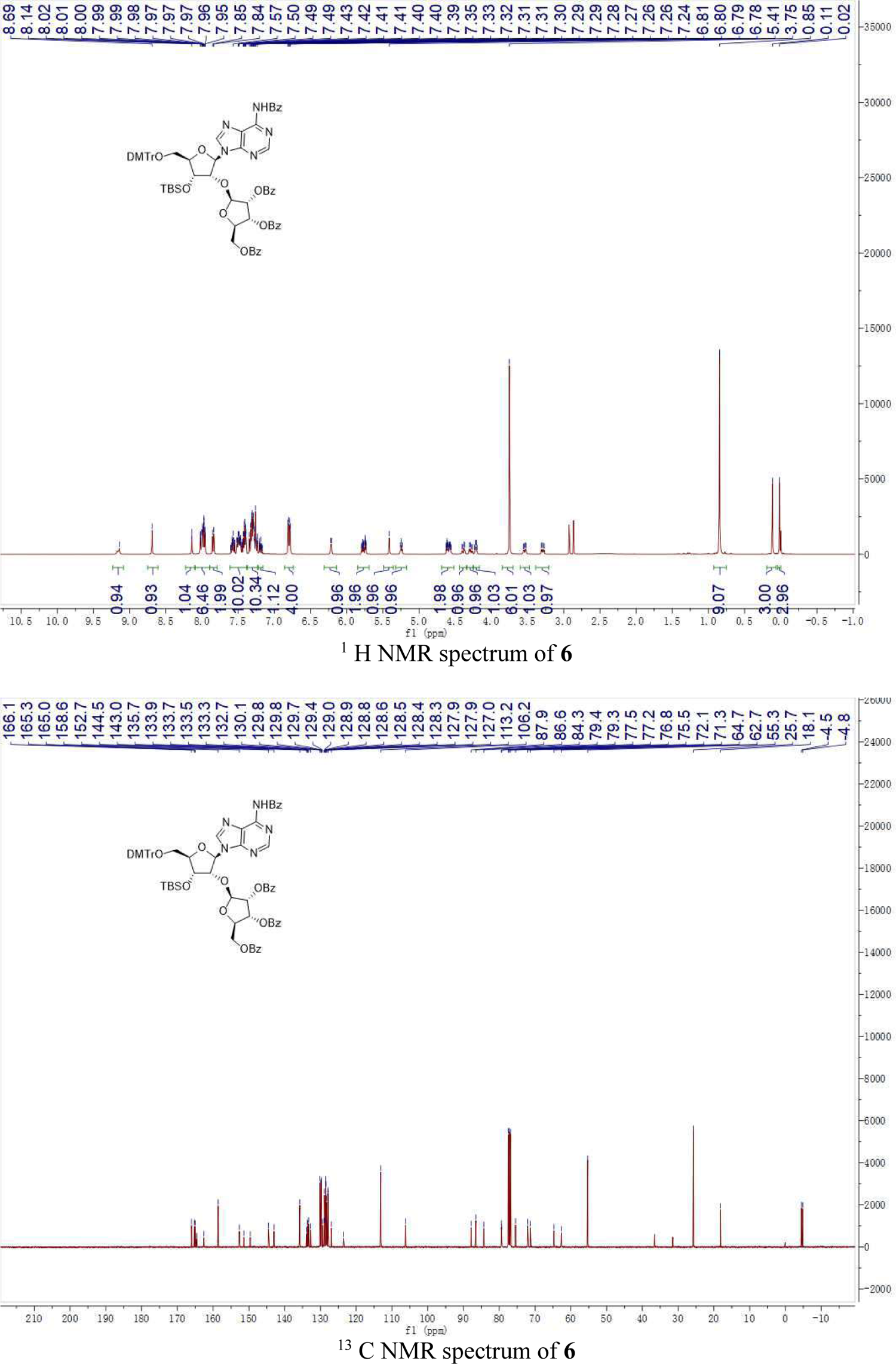
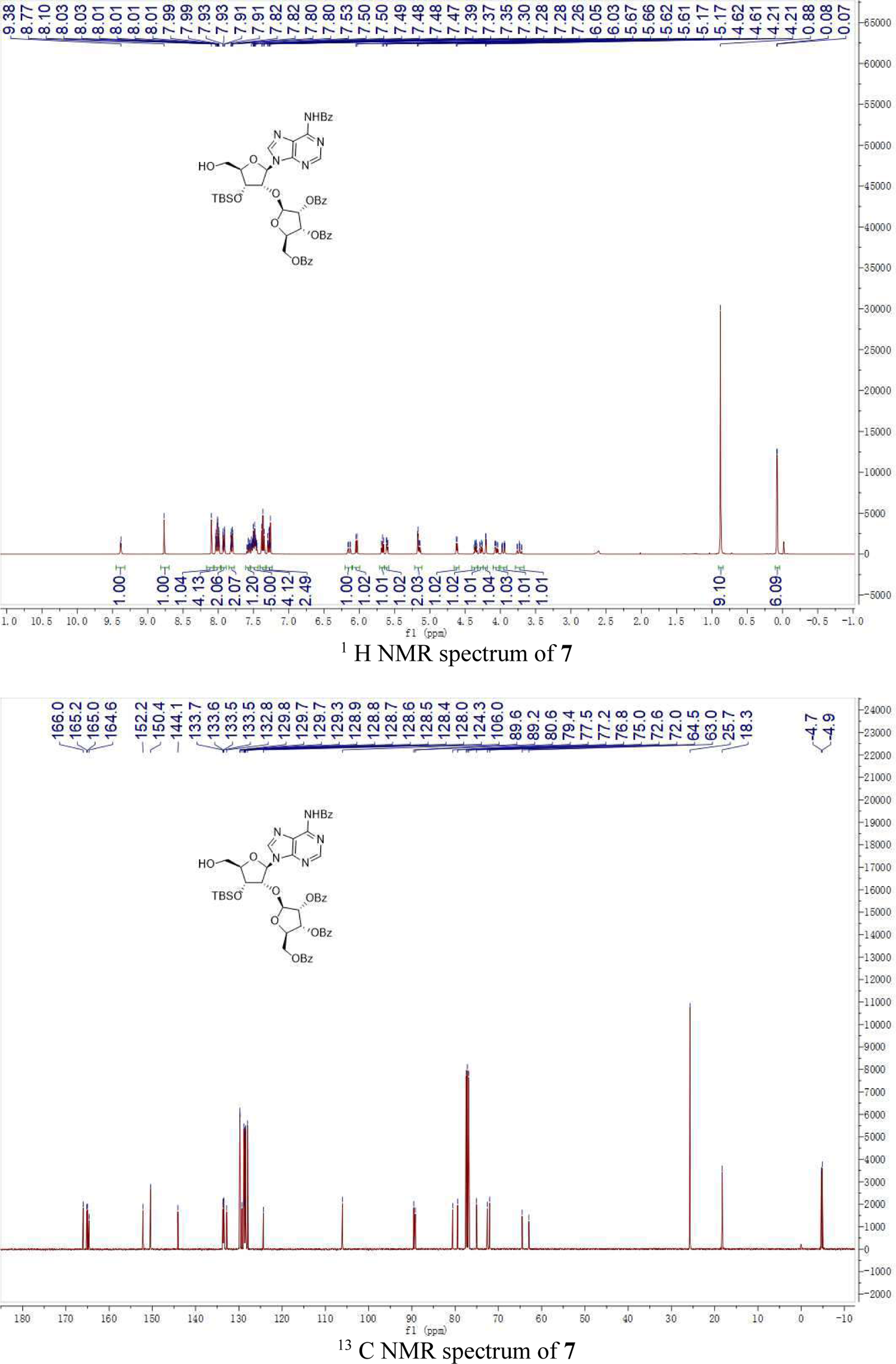
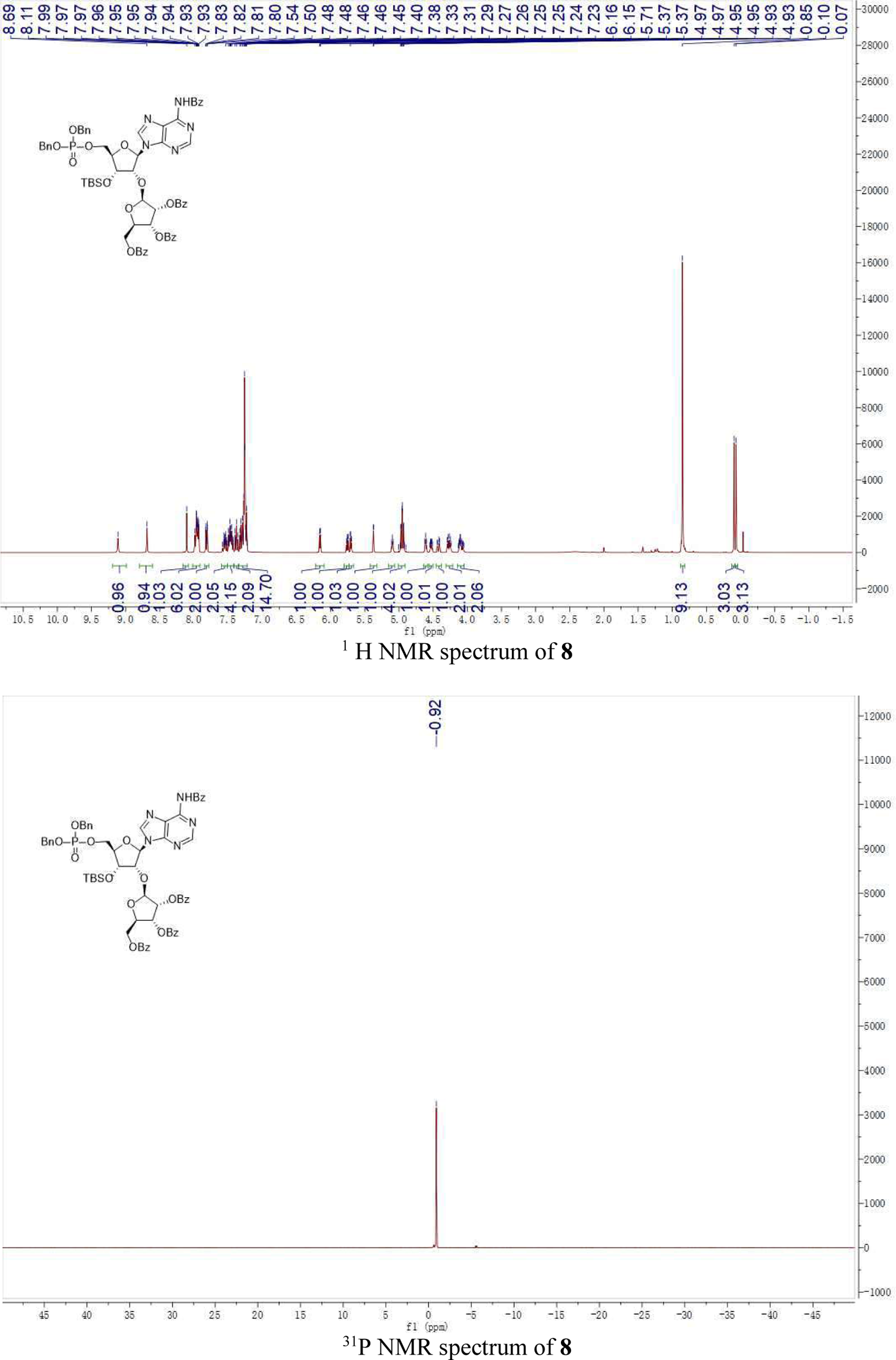
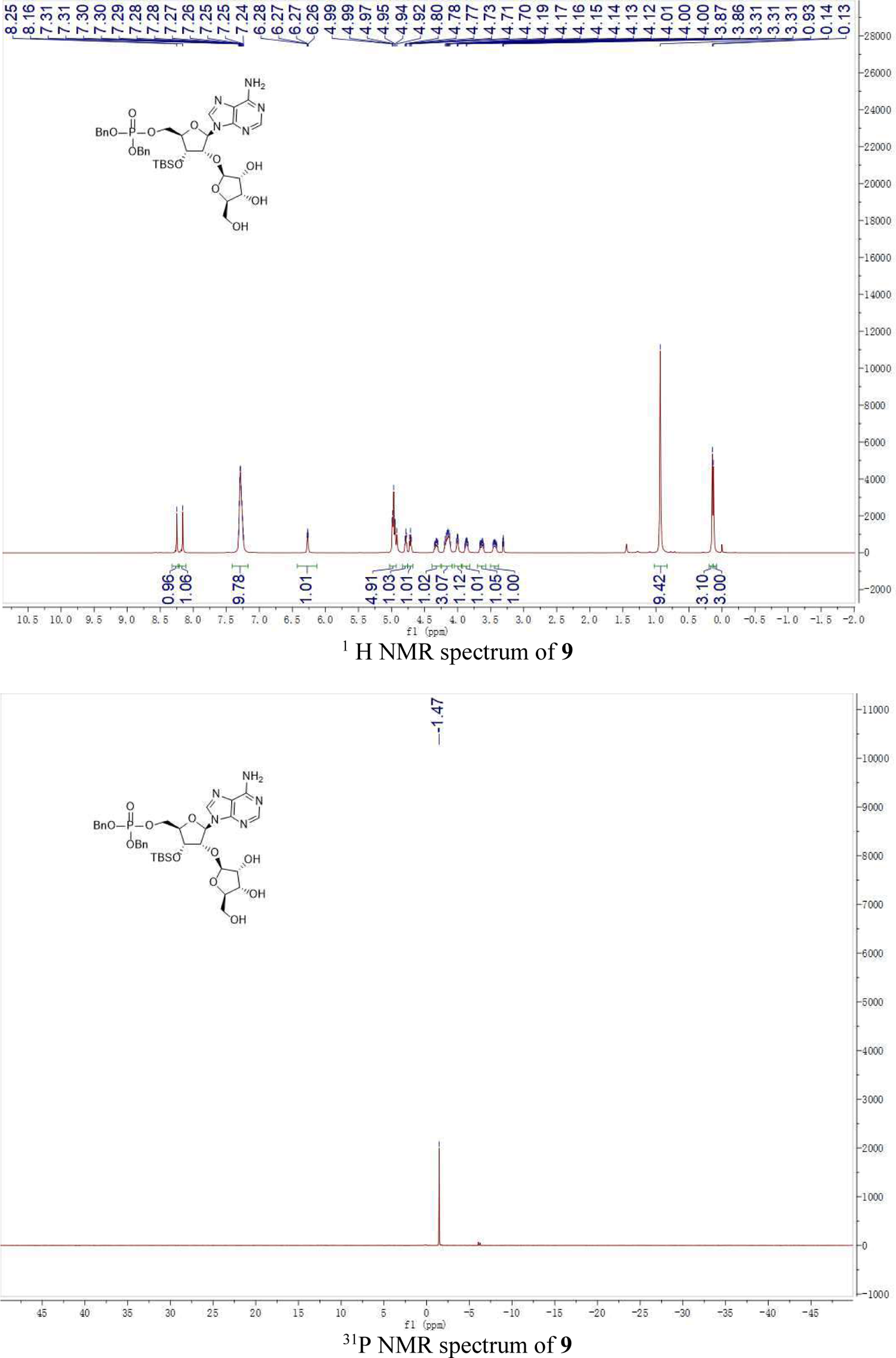
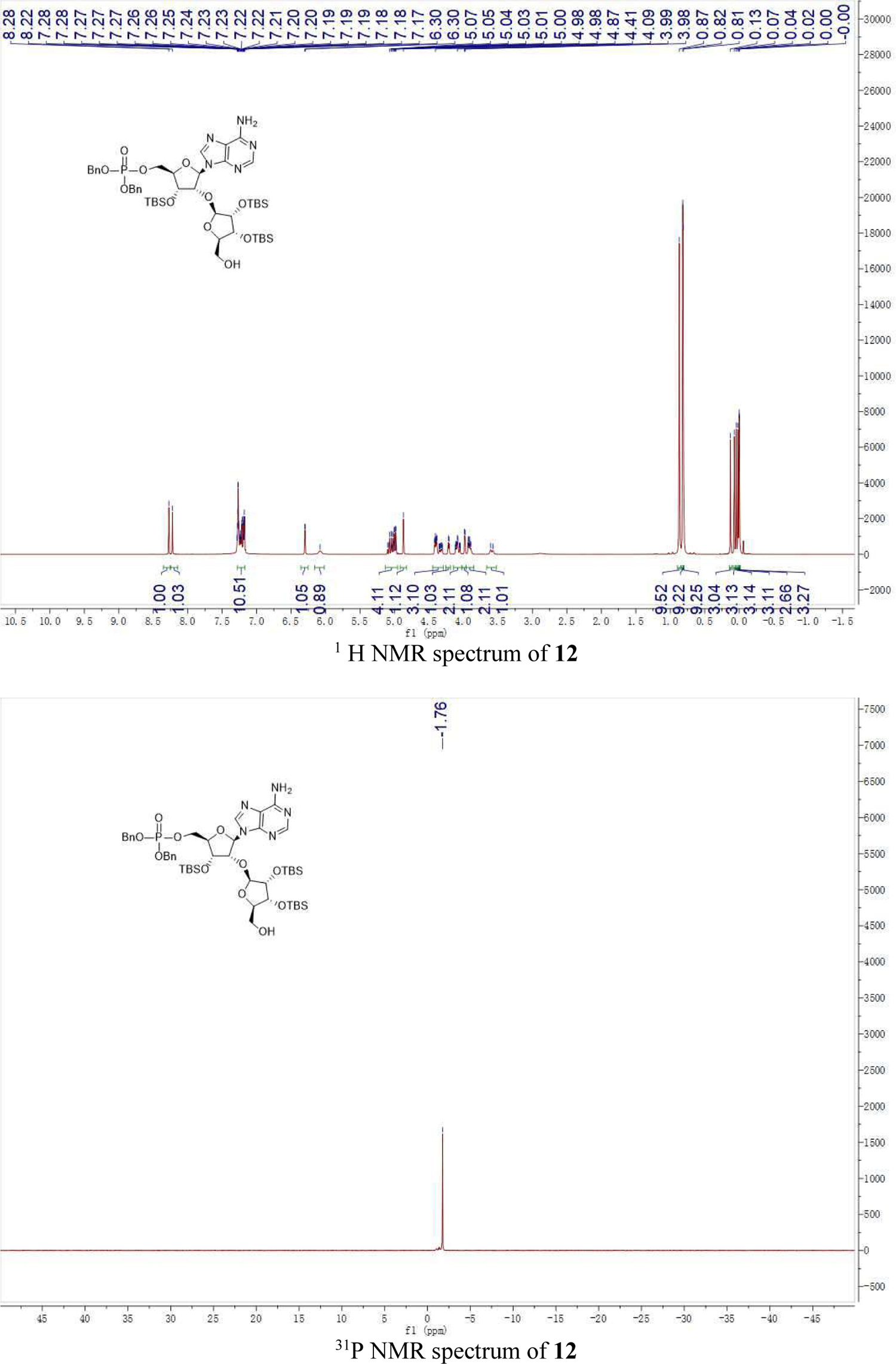
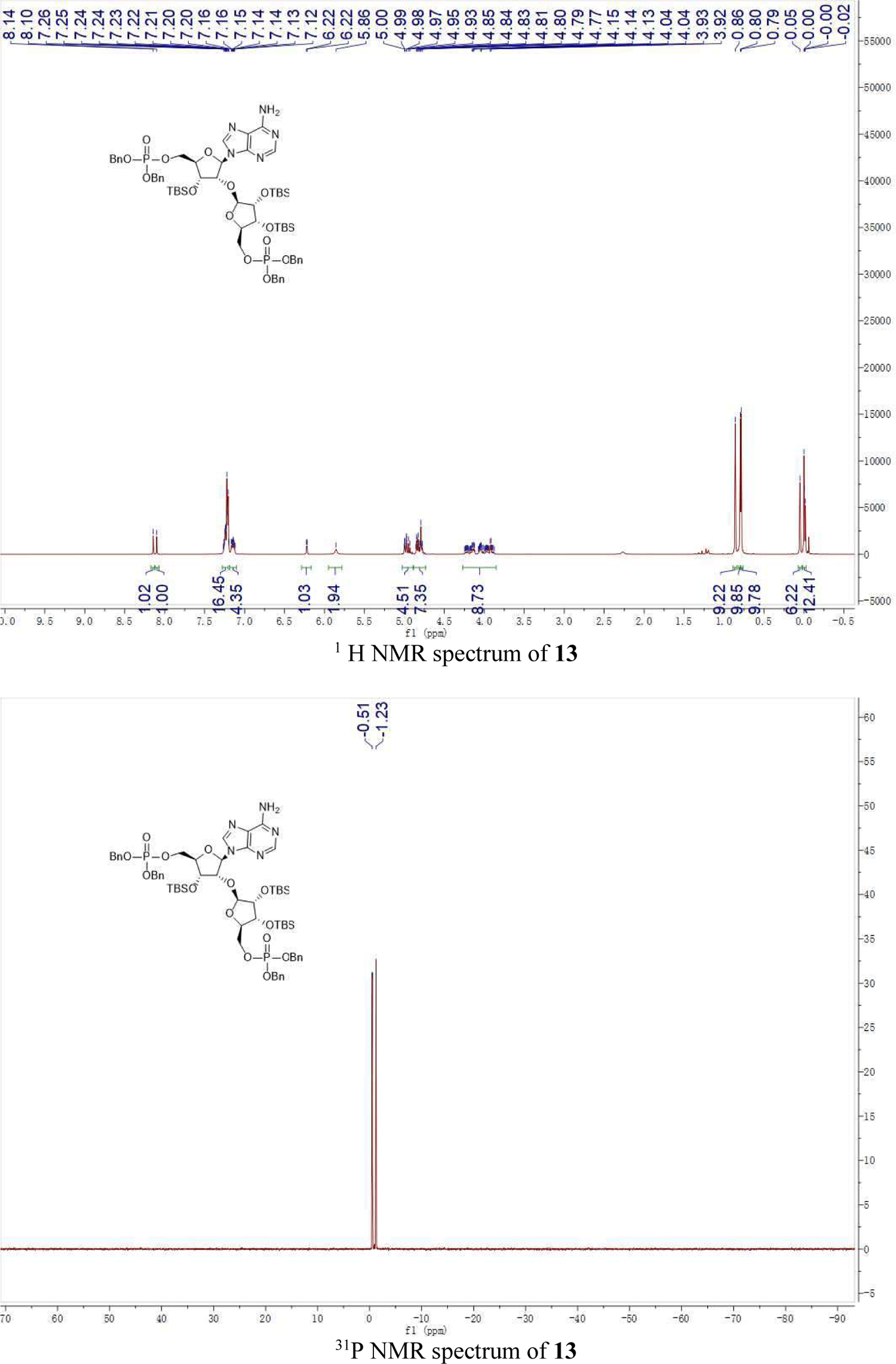
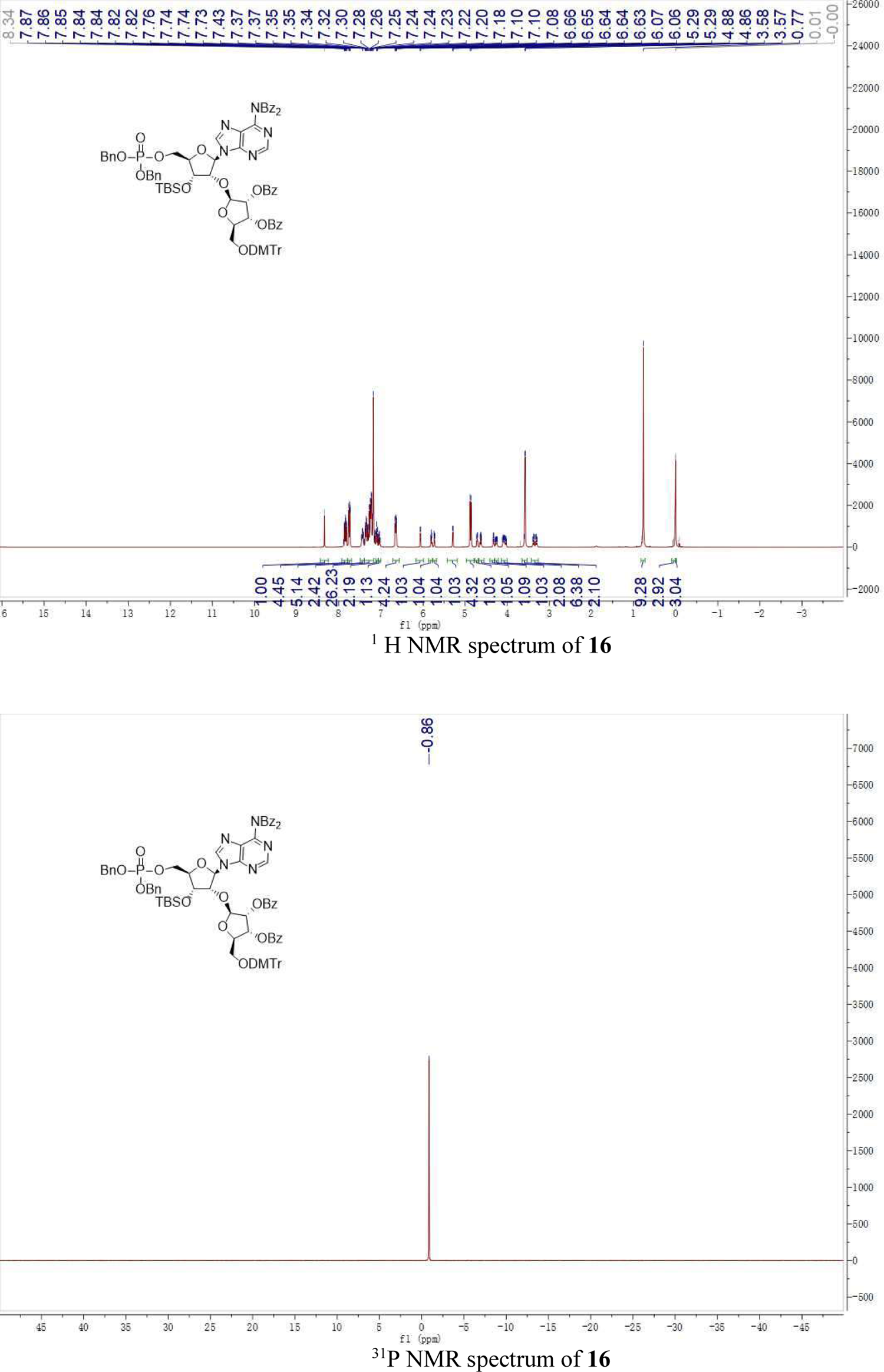
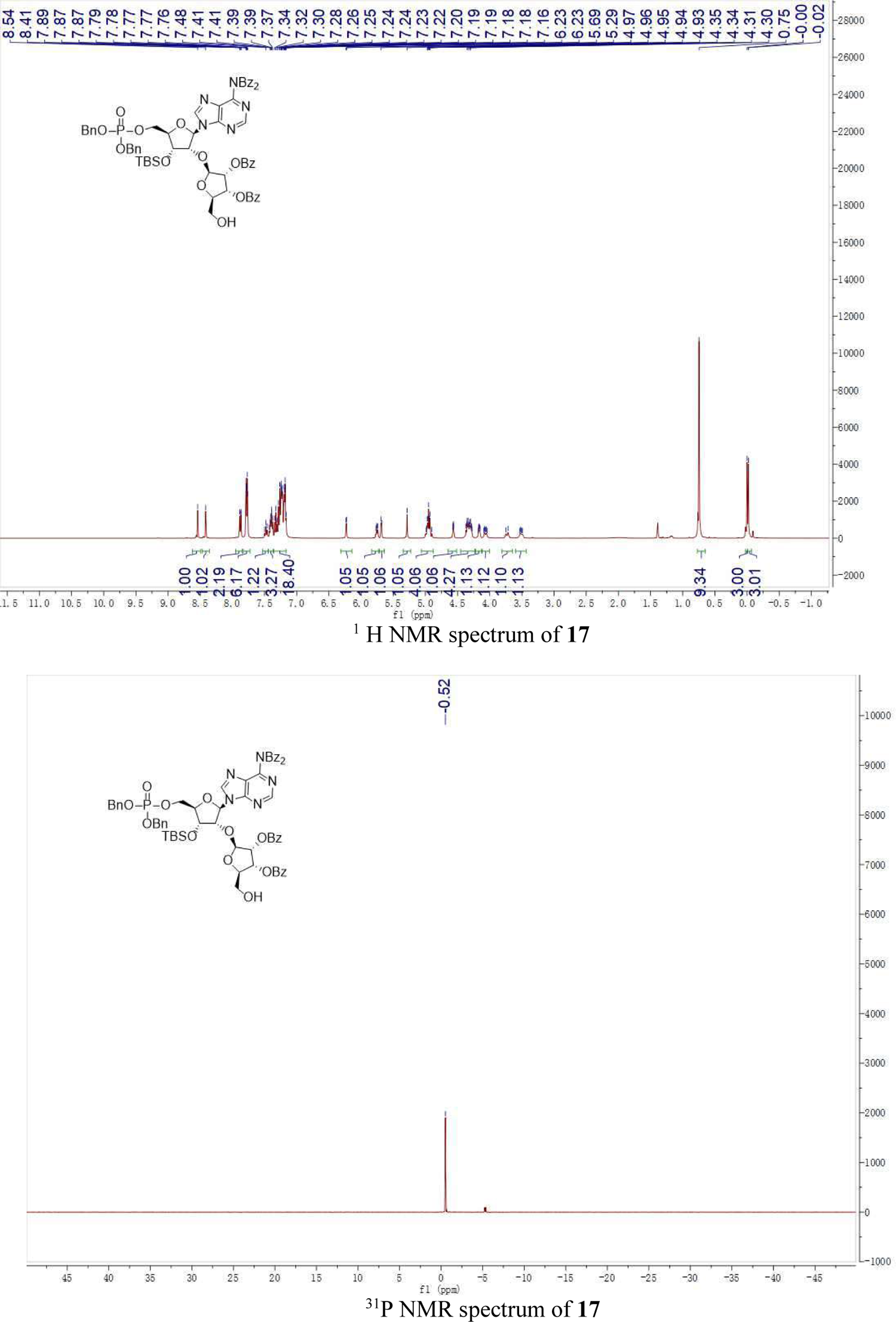
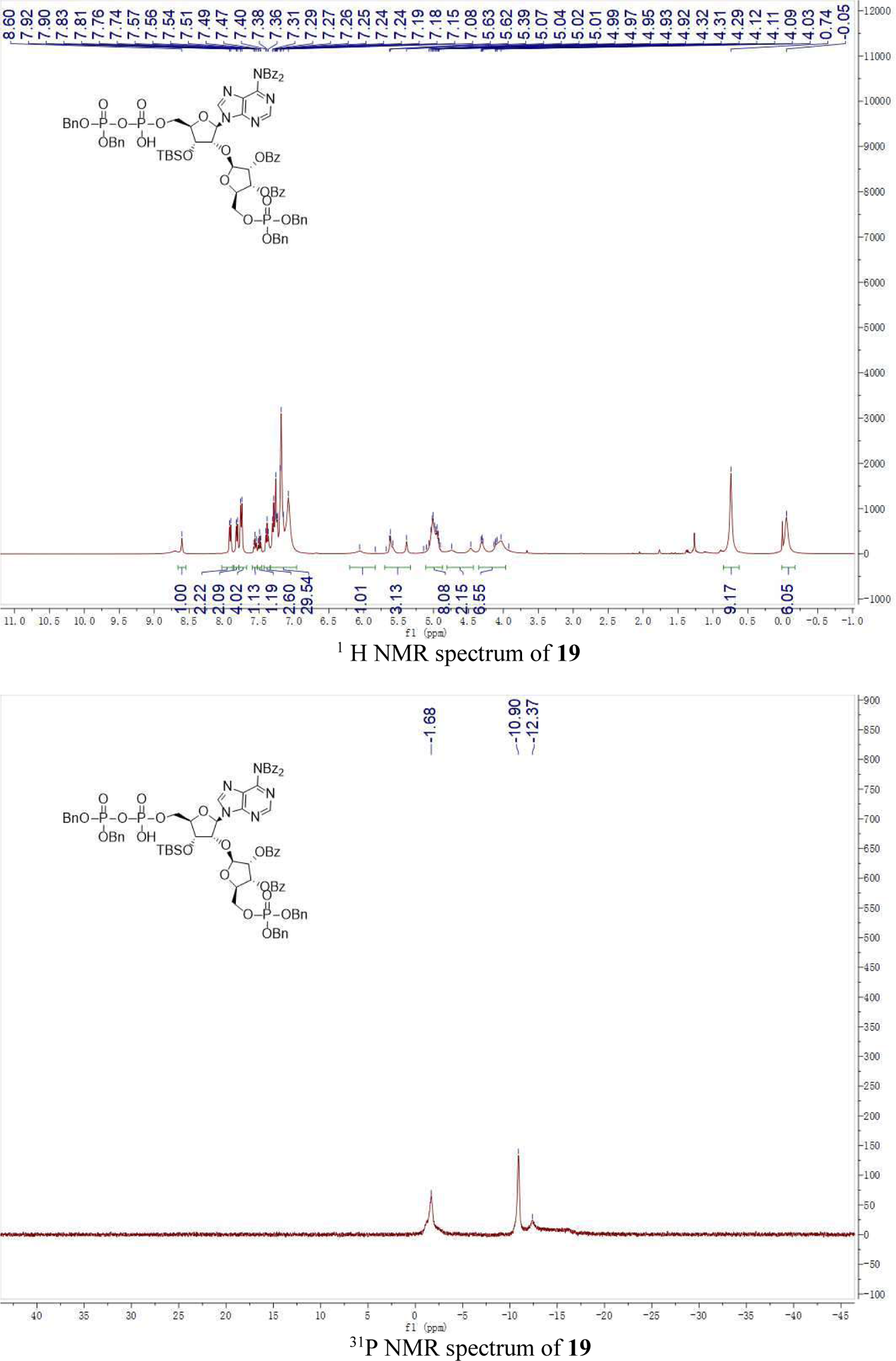

## References

1. N. Maruta et al., Structural basis of NLR activation and innate immune signalling in plants. Immunogenetics 74, 5–26 (2022).

2. B. C. Meyers, M. Morgante, R. W. Michelmore, TIR-X and TIR-NBS proteins: two new families related to disease resistance TIR-NBS-LRR proteins encoded in Arabidopsis and other plant genomes. Plant J. 32, 77–92 (2002).

3. D. Lapin, O. Johanndrees, Z. Wu, X. Li, J. E. Parker, Molecular innovations in plant TIR-based immunity signaling. Plant cell koac035 (2022).

4. J. D. Jones, R. E. Vance, J. L. Dangl, Intracellular innate immune surveillance devices in plants and animals. Science 354, aaf6395 (2016).

5. B. P. M. Ngou, P. Ding, J. D. Jones, Thirty years of resistance: Zig-zag through the plant immune system. Plant cell koac041 (2022).

6. J. Wang, J. Chai, Molecular actions of NLR immune receptors in plants and animals. Sci. China Life Sci. 63, 1303–1316 (2020).

7. Y. Xiong, Z. Han, J. Chai, Resistosome and inflammasome: platforms mediating innate immunity. Curr Opin plant Biol. 56, 47–55 (2020).

8. G. Bi et al., The ZAR1 resistosome is a calcium-permeable channel triggering plant immune signaling. Cell 184, 3528–3541 e3512 (2021).

9. S. Ma et al., Direct pathogen-induced assembly of an NLR immune receptor complex to form a holoenzyme. Science 370, eabe3069 (2020).

10. R. Martin et al., Structure of the activated ROQ1 resistosome directly recognizing the pathogen effector XopQ. Science 370, eabd9993 (2020).

11. S. Horsefield et al., NAD^+^ cleavage activity by animal and plant TIR domains in cell death pathways. Science 365, 793–799 (2019).

12. L. Wan et al., TIR domains of plant immune receptors are NAD^+^-cleaving enzymes that promote cell death. Science 365, 799–803 (2019).

13. J. M. Feehan, B. Castel, A. R. Bentham, J. D. Jones, Plant NLRs get by with a little help from their friends. Curr Opin plant Biol. 56, 99–108 (2020).

14. J. A. Dongus, J. E. Parker, EDS1 signalling: At the nexus of intracellular and surface receptor immunity. Curr Opin plant Biol. 62, 102039 (2021).

15. S. Wagner et al., Structural basis for signaling by exclusive EDS1 heteromeric complexes with SAG101 or PAD4 in plant innate immunity. Cell host Microbe 14, 619–630 (2013).

16. D. Lapin et al., A Coevolved EDS1-SAG101-NRG1 Module Mediates Cell Death Signaling by TIR-Domain Immune Receptors. Plant cell 31, 2430–2455 (2019).

17. J. Gantner, J. Ordon, C. Kretschmer, R. Guerois, J. Stuttmann, An EDS1-SAG101 Complex Is Essential for TNL-Mediated Immunity in Nicotiana benthamiana. Plant cell 31, 2456–2474 (2019).

18. D. D. Bhandari et al., An EDS1 heterodimer signalling surface enforces timely reprogramming of immunity genes in Arabidopsis. Nat. commun. 10, 772 (2019).

19. X. Sun et al., Pathogen effector recognition-dependent association of NRG1 with EDS1 and SAG101 in TNL receptor immunity. Nat. commun. 12, 3335 (2021).

20. J. A. Dongus et al., Cavity surface residues of PAD4 and SAG101 contribute to EDS1 dimer signaling specificity in plant immunity. Plant J. (2022).

21. S. M. Collier, L. P. Hamel, P. Moffett, Cell death mediated by the N-terminal domains of a unique and highly conserved class of NB-LRR protein. Mol. Plant Microbe Interact. 24, 918–931 (2011).

22. T. Qi et al., NRG1 functions downstream of EDS1 to regulate TIR-NLR-mediated plant immunity in Nicotiana benthamiana. Proc. Natl. Acad. Sci. U. S. A. 115, E10979–E10987 (2018).

23. B. Castel et al., Diverse NLR immune receptors activate defence via the RPW8-NLR NRG1. New Phytol. 222, 966–980 (2019).

24. Z. Wu et al., Differential regulation of TNL-mediated immune signaling by redundant helper CNLs. New Phytol. 222, 938–953 (2019).

25. S. C. Saile et al., Two unequally redundant “helper” immune receptor families mediate Arabidopsis thaliana intracellular “sensor” immune receptor functions. PLoS Biol. 18, e3000783 (2020).

26. Z. Wu, L. Tian, X. Liu, Y. Zhang, X. Li, TIR signal promotes interactions between lipase-like proteins and ADR1-L1 receptor and ADR1-L1 oligomerization. Plant Physiol. 187, 681–686 (2021).

27. P. Jacob et al., Plant “helper” immune receptors are Ca^2+^-permeable nonselective cation channels. Science 373, 420–425 (2021).

28. Z. Duxbury et al., Induced proximity of a TIR signaling domain on a plant-mammalian NLR chimera activates defense in plants. Proc. Natl. Acad. Sci. U.S.A. 117, 18832–18839 (2020).

29. M. R. Swiderski, D. Birker, J. D. Jones, The TIR domain of TIR-NB-LRR resistance proteins is a signaling domain involved in cell death induction. Mol. Plant Microbe Interact. 22, 157–165 (2009).

30. M. T. Nishimura et al., TIR-only protein RBA1 recognizes a pathogen effector to regulate cell death in Arabidopsis. Proc. Natl. Acad. Sci. U.S.A. 114, E2053–E2062 (2017).

31. X. Cai, Y. H. Chiu, Z. J. Chen, The cGAS-cGAMP-STING pathway of cytosolic DNA sensing and signaling. Mol. Cell 54, 289–296 (2014).

32. H. Cui et al., Antagonism of Transcription Factor MYC2 by EDS1/PAD4 Complexes Bolsters Salicylic Acid Defense in Arabidopsis Effector-Triggered Immunity. Mol. Plant 11, 1053–1066 (2018).

33. R. N. Pruitt et al., The EDS1-PAD4-ADR1 node mediates Arabidopsis pattern-triggered immunity. Nature 598, 495–499 (2021).

34. H. Tian et al., Activation of TIR signalling boosts pattern-triggered immunity. Nature 598, 500–503 (2021).

35. J. V. Gerasimenko, M. Sherwood, A. V. Tepikin, O. H. Petersen, O. V. Gerasimenko, NAADP, cADPR and IP3 all release Ca2+ from the endoplasmic reticulum and an acidic store in the secretory granule area. J. Cell Sci. 119, 226–238 (2006).

36. M. Conti, J. Beavo, Biochemistry and physiology of cyclic nucleotide phosphodiesterases: essential components in cyclic nucleotide signaling. Nat. Rev. Microbiol. 76, 481–511 (2007).

37. D. Yu et al., TIR domains of plant immune receptors are 2′,3′-cAMP/cGMP synthetases mediating cell death. *bioRxiv*, 467869 (2021).

38. Z. Otwinowski, W. Minor, Processing of X-ray diffraction data collected in oscillation mode. Methods Enzymol. 276, 307–326 (1997).

39. K. Cowtan, P. Emsley, K. S. Wilson, From crystal to structure with CCP4. Acta Crystallogr D Struct Biol. 67, 233–234 (2011).

40. P. Emsley, K. Cowtan, Coot: model-building tools for molecular graphics. Acta Crystallogr D Struct Biol. 60, 2126–2132 (2004).

41. J. Lei, J. Frank, Automated acquisition of cryo-electron micrographs for single particle reconstruction on an FEI Tecnai electron microscope. J. Struct. Biol 150, 69–80 (2005).

42. S. Q. Zheng et al., MotionCor2: anisotropic correction of beam-induced motion for improved cryo-electron microscopy. Nat. methods 14, 331–332 (2017).

43. J. A. Mindell, N. Grigorieff, Accurate determination of local defocus and specimen tilt in electron microscopy. J. Struct. Biol. 142, 334–347 (2003).

44. S. H. Scheres, RELION: implementation of a Bayesian approach to cryo-EM structure determination. J. Struct. Biol. 180, 519–530 (2012).

45. S. H. Scheres, Processing of Structurally Heterogeneous Cryo-EM Data in RELION. Methods Enzymol. 579, 125–157 (2016).

46. S. H. Scheres, A Bayesian view on cryo-EM structure determination. J. Mol. Biol. 415, 406–418 (2012).

47. P. B. Rosenthal, R. Henderson, Optimal determination of particle orientation, absolute hand, and contrast loss in single-particle electron cryomicroscopy. J. Mol. Biol. 333, 721–745 (2003).

48. A. Kucukelbir, F. J. Sigworth, H. D. Tagare, Quantifying the local resolution of cryo-EM density maps. Nat. methods 11, 63–65 (2014).

49. E. F. Pettersen et al., UCSF Chimera--a visualization system for exploratory research and analysis. J. Comput. Chem. 25, 1605–1612 (2004).

50. P. Emsley, B. Lohkamp, W. G. Scott, K. Cowtan, Features and development of Coot. Acta Crystallogr D Struct Biol. 66, 486–501 (2010).

51. P. D. Adams et al., PHENIX: a comprehensive Python-based system for macromolecular structure solution. Acta Crystallogr D Struct Biol. 66, 213–221 (2010).

